# Mapping cortical excitability in the human dorsolateral prefrontal cortex

**DOI:** 10.1101/2023.01.20.524867

**Authors:** Juha Gogulski, Christopher C. Cline, Jessica M. Ross, Jade Truong, Manjima Sarkar, Sara Parmigiani, Corey J. Keller

**Affiliations:** Department of Psychiatry & Behavioral Sciences, Stanford University Medical Center, Stanford, CA, 94305, USA; Wu Tsai Neuroscience Institute, Stanford University, Stanford, CA, USA; Department of Clinical Neurophysiology, HUS Diagnostic Center, Clinical Neurosciences, Helsinki University Hospital and University of Helsinki, Helsinki, FI-00029 HUS, Finland; Veterans Affairs Palo Alto Healthcare System, and the Sierra Pacific Mental Illness, Research, Education, and Clinical Center (MIRECC), Palo Alto, CA, 94394, USA

**Keywords:** TMS-EEG, Transcranial magnetic stimulation (TMS), electroencephalography (EEG), dorsolateral prefrontal cortex (dlPFC), TMS-evoked potentials (TEPs)

## Abstract

**Objective:** To characterize early TEPs anatomically and temporally (20-50 ms) close to the TMS pulse (EL-TEPs), as well as associated muscle artifacts (<20 ms), across the dlPFC. We hypothesized that TMS location and angle influence EL-TEPs, and that EL-TEP amplitude is inversely related to muscle artifact. Additionally, we sought to determine an optimal group-level TMS target and angle, while investigating the potential benefits of a personalized approach.

**Methods:** In 16 healthy participants, we applied single-pulse TMS to six targets within the dlPFC at two coil angles and measured EEG responses.

**Results:** Stimulation location significantly influenced EL-TEPs, with posterior and medial targets yielding larger EL-TEPs. Regions with high EL-TEP amplitude had less muscle artifact, and vice versa. The best group-level target yielded 102% larger EL-TEP responses compared to other dlPFC targets. Optimal dlPFC target differed across subjects, suggesting that a personalized targeting approach might boost the EL-TEP by an additional 36%.

**Significance:** Early local TMS-evoked potentials (EL-TEPs) can be probed without significant muscle-related confounds in posterior-medial regions of the dlPFC. The identification of an optimal group-level target and the potential for further refinement through personalized targeting hold significant implications for optimizing depression treatment protocols.

**Highlights:**

- Early local TMS-evoked potentials (EL-TEPs) varied significantly across the dlPFC as a function of TMS target.
- TMS targets with less muscle artifact had significantly larger EL-TEPs.
- Selection of a postero-medial target increased EL-TEPs by 102% compared to anterior targets.

## 1. Introduction

Transcranial magnetic stimulation (TMS) to the dorsolateral prefrontal cortex (dlPFC) is a safe and effective treatment for medication-resistant depression (Mutz et al., 2018). Despite widespread clinical adoption, however, the neural effects of TMS treatment remain unknown. As changes in dlPFC excitability are thought to underlie the clinical effects of TMS (Eshel et al., 2020; Voineskos et al., 2021), brain-based metrics that track cortical excitability with high signal quality are paramount to improve our understanding of the neural effects of TMS treatment and guide future treatment optimization.

Causal noninvasive measures of local cortical excitability exist but to date have not been well-characterized across different dlPFC treatment targets (Ferrarelli and Phillips, 2021). Single pulses of TMS paired with electroencephalography (TMS-EEG) provide a noninvasive, causal readout of neural activity with millisecond resolution sufficient to capture treatment-related changes in cortical excitability (Voineskos et al., 2021). Moreover, single pulse TMS-evoked EEG potentials (TEPs (Lioumis et al., 2009)) in areas outside of the dlPFC have been used to monitor sleep stages (Bergmann et al., 2012; Massimini et al., 2007), measure the neural effects to electroconvulsive therapy (Casarotto et al., 2013), and differentiate neurovegetative states from coma (Gosseries et al., 2014). Despite these use cases, our understanding of TEPs evoked by TMS targeting the dlPFC remains limited. This is in part due to the large TMS-induced muscle artifacts, which have been difficult to remove and often confound TEPs (Mutanen et al., 2013). Furthermore, only a few TEP studies have directly compared TMS-induced muscle artifacts across the dlPFC. Studies have explored this TEP / artifact relationship both at one dlPFC site (Eshel et al., 2020; Kerwin et al., 2018; Rogasch et al., 2014, 2013) as well as in more medial prefrontal cortex (Casarotto et al., 2022). However, to date characterization of early local TEPs (EL-TEPs) and artifacts across clinically relevant sub-regions of the dlPFC is lacking. Addressing these gaps will enhance our understanding of which clinical TMS targets can be effectively monitored using TMS-EEG, as well as the potential benefits of target personalization to enhance the signal quality of TEPs.

Here we sought to map the distribution of EL-TEPs and TMS-induced muscle artifacts across the dlPFC. We focused on the early (20-50 ms) TEP in the left dlPFC as it represents the earliest noninvasively measurable neural response to TMS (Casarotto et al., 2022), appears to reflect local cortical excitability (Darmani et al., 2019), and avoids later responses (>100 ms) that at least in part reflect off-target sensory responses (Biabani et al., 2019; Ross et al., 2022a, 2022b). In 16 healthy participants, we applied single-pulse TMS to six clinically-relevant dlPFC locations and two coil orientations. For each target / angle combination, we computed both the EL-TEP and the TMS-induced muscle artifact. Based on the anatomical distribution of facial muscles (Mäki and Ilmoniemi, 2011), which directly relates to the strength of evoked muscle artifact, we hypothesized that 1) posterior-medial regions would induce the smallest artifacts and largest EL-TEPs, and 2) an individualized approach to minimize artifact may yield larger EL-TEPs and motivate personalized targeting for future clinical applications. Across dlPFC targets, we observed both EL-TEPs and artifacts were sensitive to target location, with targets evoking larger artifacts generally associated with smaller EL-TEPs. The group-best target / angle location yielded EL-TEPs that were more than twice the size of other targets. Individualized target / angle combination produced larger EL-TEPs than the group-best combination, suggesting that a personalized targeting strategy may be effective. Taken together, these findings inform our ability to causally measure cortical excitability across the dlPFC and suggest a personalized approach may be beneficial. This work has potential application in treatment monitoring and biomarker-driven treatment optimization.

### 2. Methods

### 2.1. Participants

22 healthy participants (23-64 years old, mean=39.45, SD=13.00, seven females) were recruited into the study and provided written informed consent under a protocol approved by the Stanford University Institutional Review Board. All participants completed an online screening questionnaire with inclusion criteria as follows: aged 18-65, fluent in English, able to travel to study site, and fully vaccinated against COVID-19. Participants completed the Quick Inventory of Depressive Symptomatology (16-item, QIDS) self-report questionnaire and were excluded if they scored 11 or higher, indicating moderate or more severe depression (Rush et al., 2003). Additional criteria for exclusion included a lifetime history of psychiatric or neurological disorder, substance or alcohol abuse / dependence in the past month, recent heart attack (<3 months), pregnancy, the presence of any contraindications for TMS (Rossi et al., 2011, 2009), or usage of psychotropic medications. All eligible participants were scheduled for two study visits on separate days to first obtain an MRI and then complete an TMS-EEG session. All enrolled participants completed an MRI pre-examination screening form provided by Richard M. Lucas Center for Imaging at Stanford University. Six subjects were excluded from the study: three before the experimental session, one for TMS intolerability assessed by delivering test pulses before the experiment, one for lower back pain, and one for technical difficulties. All remaining 16 subjects (24-64 years old, mean=38.81, SD=13.39, three females) were included in the analyses.

### 2.2. Transcranial Magnetic Stimulation

Single-pulse TMS was delivered using a MagVenture Cool-B65 A/P figure-of-eight coil with a MagPro X100 stimulator (MagVenture, Denmark). A thin (0.5 mm) foam pad was attached to the TMS coil to minimize electrode movement and bone-conducted auditory artifact (ter Braack et al., 2015). The TMS coil was held and positioned automatically by a robotic arm (TMS Cobot, Axilum Robotics, France). MRI-based neuronavigation (Localite TMS Navigator, Localite, Germany) was used to align the coil to pre-planned individualized coil orientations for each stimulation target (see section 2.2.3). To estimate resting motor threshold (rMT), biphasic single pulses of TMS were delivered to the hand region of the left motor cortex with the coil held tangentially to the scalp and at 45° from midline. The optimal motor hotspot was defined as the coil position from which TMS produced the largest and most consistent visible twitch in relaxed right first dorsal interosseous muscle (FDI; (Badran et al., 2019)). rMT was determined to be the minimum intensity that elicited a visible twitch in relaxed FDI in ≥ 5 out of 10 pulses.

#### 2.2.1 dlPFC Target Selection

For each subject, single-pulse TMS was delivered to six different targets within the left dlPFC (Table 1). The number of targets and spatial locations were chosen to balance the following: sampling of the anterior / posterior and medial / lateral planes in the middle frontal gyrus and sampling of the many of the utilized clinical dlPFC targets for depression (Cash et al., 2021), as shown in Fig. S1. While higher density sampling is ideal for full characterization, this number of targets allowed for sampling of much of the dlPFC and to test two different angles within a reasonable session duration for the participants. The MNI coordinates for these six targets are outlined in Table 1. The *t1* target is close to the commonly used 5 cm average clinical target (Fox et al., 2012). The *t2* target corresponds to the EEG F3 site of Herwig et al., 2003, as presented by Cash et al., 2021. The *t3* target is located between the BA9 definition by Rajkowska and Goldman-Rakic, 1995 and a treatment site with activity negatively correlated to anterior subgenual cingulate cortex (sgACC (Fox et al., 2012)). The *t4* target corresponds to a treatment site with activity that is most negatively correlated to the sgACC, as described by Fox et al., 2012. The *t5* target is close to a TMS target that produced clinical response in the study by Herbsman et al., 2009. The *t6* target is close to the 5.5 cm average target of Weigand et al., 2018. Euclidean distances between the stimulation sites of the current study are shown in Table S1, and the distances between the current targets and clinical targets are depicted in Fig. S2. The six targets were registered from MNI space to each individual brain using a non-linear inverse transformation (see 2.3.).

**Table 1.** MNI coordinates of the dlPFC stimulation targets. . BA: Brodmann area (according to MRICron Brodmann atlas (https://www.nitrc.org/projects/mricron). HCPex: Brodmann areas according to HCPex atlas (Huang et al., 2022). *Locations of the clinical targets were extracted from a summary of Cash et al., 2021. **Target t4 had the same MNI coordinates as a treatment site with activity that is most negatively correlated to the sgACC, as described by Fox et al., 2012.

| Stimulation site | x | y | z | BA | HCPex | Closest clinical target* |
| --- | --- | --- | --- | --- | --- | --- |
| t1 | -37 | 12 | 57 | 9 | 8av | 5 cm average (Fox *et al.*, 2012) |
| t2 | -37 | 23 | 47 | 9 | 8av | EEG F3 site (Herwig *et al.*, 2003) |
| t3 | -38 | 33 | 36 | 46 | 46 | Optimal FC Group Target #1 (Fox *et al.*, 2012) |
| t4 | -38 | 44 | 26 | 46 | 46 | Optimal FC Group Target #2 (Fox *et al.*, 2012)** |
| t5 | -42 | 20 | 42 | 44 (/9) | NaN/8Av | Clinical responders (Herbsman *et al.*, 2009) |
| t6 | -32 | 26 | 52 | 8 (/9) | 8Av | 5.5cm average (Weigand *et al.*, 2018) |

#### 2.2.2. Experimental Design

The order of target / angle conditions was pseudorandomized within the session. 150 single-pulse TMS trials were collected for each experiment block. For a subset of conditions (all targets at 45°), two blocks of single pulse data were collected. For these conditions, the TEP amplitudes from each set of 150 single pulse blocks were averaged together. For each condition, single TMS pulses were delivered every 1.5 s, jittered +/- 25%. In the Supplementary Material (Fig. S3) we show that this relatively short interval between the TMS pulses does not elicit plasticity effects. To maintain levels of attention, subjects were shown neutral images on a screen and were instructed to keep eyes open during stimulation. Subjects were monitored throughout the experiment.

### 2.3. Stimulation target and intensity individualization

Before each experimental session, electric fields (E-fields) induced by TMS were computationally modeled using SimNIBS (Nielsen et al., 2018; Thielscher et al., 2015) (v3.2.5). Subject-specific head models were generated from T1 and T2 MRIs using *headreco*. Deformation fields providing a non-linear spatial mapping from MNI coordinates to the brains of individual participants were generated using SPM12 (https://www.fil.ion.ucl.ac.uk/spm/). T1 MRIs were segmented and inverse deformation fields were created for each individual. MNI target coordinates were warped to individual space using the inverse deformation field (Fig. 1A). Given a target coordinate in individual brain space, 45° and 90° coil orientations were generated based on positioning the coil perpendicular to the nearest point on the scalp, with an additional offset to account for EEG electrode and coil foam thickness. Using a custom dipole model of the TMS coil, induced E-fields were then predicted for each coil orientation (Fig. 1B).

**Figure 1.**
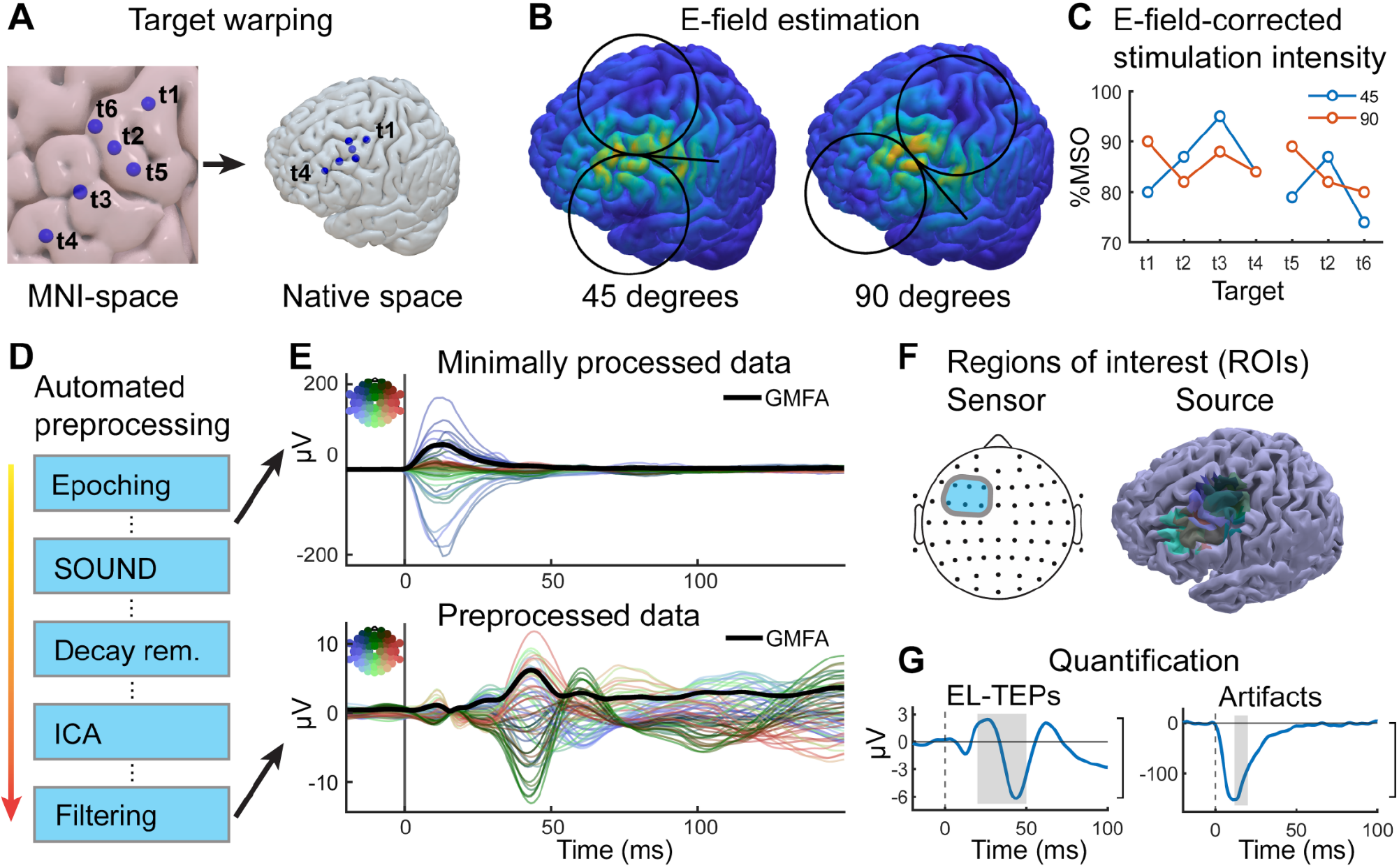
Study schematic. A) Targets for single pulse TMS within the dlPFC in MNI and native space. Stimulation targets were warped to individual space for each subject. B) Before the TMS-EEG session, maximal E-field was calculated for each target / angle combination. The stimulation intensity was adjusted for each target / angle combination so that the maximal induced E-field was the same in the dlPFC as compared to the intensity of 120% rMT in the primary motor cortex. Shown are E-fields for one target with 45° (left) and 90° (right) angles in a representative subject. C) Adjusted stimulation intensities for each target and angle in a representative subject. D) The data were preprocessed using a fully automatized preprocessing pipeline, described in detail in (Cline et al., 2021). For each condition, artifact amplitude was quantified before artifact removal (SOUND). E) Example butterfly plots (one target / angle condition, 150 trials, N=1 subject). F) For TEP analyses in sensor space, an ROI consisting of six prefrontal electrodes (F5, F3, F1, FC5, FC3, FC1) was used (left). For TEP analyses in source space, each target has an individual, 20 mm spherical ROI, positioned around the stimulation location. Source-space ROIs are shown for a representative subject (right). G) Peak-to-peak amplitudes of EL-TEPs (left) and maximal values of artifacts (right) were quantified. Artifacts were quantified in sensor space. One condition in sensor space is shown for a representative subject. Brackets denote the amplitude of EL-TEP and artifact.

Maximal E-field magnitudes on the cortical surface were calculated before each session (after rejecting the top 0.01% of values to remove outliers) to obtain an estimate of stimulation intensity at the cortical surface (Fig 1B and Fig. S4). During the experiments, the stimulation intensities (Fig. 1C) were set based on the experimentally-determined motor threshold multiplied by the ratio between maximal predicted E-field magnitude at the site of stimulation and at the motor hotspot, similar to the APEX MT approach (Caulfield et al., 2021). In theory, this procedure results in a consistent stimulation intensity at the brain across all stimulation targets, accounting for variation in anatomical parameters such as skull thickness across targets. Target stimulation intensities for prefrontal targets were set to produce, at each target, 110% of the E-field magnitude that at M1 corresponded to rMT. If the calculated stimulation intensity exceeded 100% maximal stimulator output (MSO) for any target, intensities of all conditions were scaled down in such a way that the largest intensity would not exceed 100% MSO (8/16 subjects). Moreover, if the subject did not tolerate stimulation of one of the conditions, the intensities of all conditions were lowered (7/16 subjects). Tolerability of each target / angle combination was tested before the actual experiment. Across all stimulation targets and subjects, the TMS intensity varied from 45 to 100% MSO (mean 76% MSO, SEM ±0.83; mean %rMT 110, SEM ±0.98; Fig. S5-6, Table S2). To test whether slight TMS intensity differences at each target, due to E-field modeling, could account for differences across targets observed, we calculated the correlation of EL-TEP amplitude and stimulation intensity for each individual. There was no association between stimulation intensity and TEP amplitude (mean *R*=0.22, Fig. S7).

### 2.4. Electroencephalography

EEG was recorded using a 64-channel TMS-compatible amplifier (ActiCHamp Plus, Brain Products GmbH, Germany) with a 25 kHz sampling rate. Slim, freely rotatable, active electrodes (actiCAP slim, Brain Products GmbH, Germany) were used in a standard montage labeled according to the extended 10-20 international system. EEG data were online referenced to the ‘Cz’ electrode and recorded using BrainVision Recorder software (Brain Products GmbH, Germany). Impedance at all electrodes were generally kept below 5 kΩ and monitored at regular intervals throughout each session. Noise masking and earmuffs, as described in detail previously (Ross et al., 2022b), were applied to reduce off-target sensory effects.

#### 2.4.1. Preprocessing of TMS-EEG data

TMS-EEG preprocessing was performed with version 2 of the fully automated AARATEP pipeline (Cline et al., 2021) (Fig. 1D). Epochs were extracted from 800 ms before to 1000 ms after each TMS pulse. Data between 2 ms before to 12 ms after each pulse were replaced with values interpolated by autoregressive extrapolation and sigmoidal-weighted blending between pre-stim and post-stim time periods (Cline et al., 2021), downsampled to 1 kHz, and baseline-corrected based on mean values between 500 to 10 ms before the pulse. Epochs were then high-pass filtered above 1 Hz with a modified filtering approach to reduce the spread of pulse-related artifact into baseline time periods; this filtering stage involved separate autoregressive extrapolation forward from the pre-stimulation period and backward from the post-stimulation period, piecewise high-pass filtering on these two fitted plus extrapolated signals, sigmoidal-weighted blending, and then applying the difference in these signals as a result of filtering to the original non-extrapolated data. See (Cline et al., 2021) and code at github.com/chriscline/AARATEPPipeline for details of this modified filtering procedure. Bad channels were identified based on a Wiener noise estimate threshold of 10 (Mutanen et al., 2016), and replaced with spatially interpolated values. Eye blink artifacts were attenuated by a dedicated round of independent component analysis (ICA) and eye-specific component labeling and rejection using ICLabel (Pion-Tonachini et al., 2019), a modification from the original AARATEP pipeline introduced in version 2. An average of 1.7 +/- 0.7 components were rejected at this stage. Various non-neuronal noise sources were attenuated with SOUND (Mutanen et al., 2018). Decay artifacts were further reduced via a specialized decay fitting and removal procedure, involving estimation of the dominant low-latency artifact topography, exponential fitting on the trial-averaged artifact time course, and constrained removal of per-trial artifact signals based on the decay fit, minimizing removal of distinct, later latency TEP components even if their topography was similar to the decay artifact. See (Cline et al., 2021) and code at github.com/chriscline/AARATEPPipeline for details of this decay removal procedure. Line noise was attenuated with a bandstop filter between 58-62 Hz. Additional artifacts were attenuated with a second stage of ICA and ICLabel labeling and rejection, with rejection criteria targeted at removing any clearly non-neural signals (see (Cline et al., 2021) for all data deletion criteria). 60.9% +/- 10.5% of components were rejected, accounting for 30.8% +/- 21.0% of variance in the signals at this stage. Data were again interpolated between -2 and 12 ms with autoregressive extrapolation and blending, low-pass filtered below 100 Hz, and average re-referenced. For complete details of the pipeline implementation, see (Cline et al., 2021) and source code at github.com/chriscline/AARATEPPipeline.

#### 2.4.2. Extraction of EL-TEP and artifact amplitudes in sensor space

Our goal was to characterize the local TMS-evoked neural responses and muscle artifacts across the dlPFC. To do so, we computed two measures: 1) the *EL-TEP*, defined as the peak-to-peak amplitude (subtraction of the signal minimum from the signal maximum) within 20 to 50 ms after the TMS pulse; and 2) *early artifact*, induced primarily by local muscle activation (Mäki and Ilmoniemi, 2011), and computed as the maximal amplitude from 12-20 ms from the minimally processed (after epoching and baseline correction) data, in dBμV (Fig. 1G). We focused on the early time window of the TEP, which encompasses the ‘P20’ and the ‘N40’ complexes (Kähkönen et al., 2005; Lioumis et al., 2009; Tremblay et al., 2019), because 1) there is evidence for the neural basis of the EL-TEP components (Huang et al., 2019; Keller et al., 2018); 2) in other brain regions it has been well-characterized by other groups (Ahn and Fröhlich, 2021; Hill et al., 2016; Kähkönen et al., 2005; Lioumis et al., 2009; Tremblay et al., 2019); and 3) off-target sensory confounds are more prominent in the later (>100 ms) components (Biabani et al., 2019; Conde et al., 2019; Freedberg et al., 2020, p.; Rocchi et al., 2021; Ross et al., 2022a, 2022b). Because the timing of the P20 and N40 are variable across locations and subjects, we simplified and quantified the peak-to-peak amplitude across the time window that captures both peaks. For both *EL-TEP* and *early artifact* measures, a region of interest (ROI) consisting of six left prefrontal electrodes (F5, F3, F1, FC5, FC3, FC1) was used (Fig. 1F). These electrodes were chosen to broadly cover the dlPFC. To make sure that our *a priori* chosen ROI did not skew the results towards targets closer to the center of the ROI, we also computed the results using two ‘shifted ROIs’ (Fig. S8). Trial-averaged TEPs were calculated with a 10% trimmed mean to prevent a small number of extreme trials from dominating results.

#### 2.4.3. Source imaging

Subject-specific differences in gyral anatomy can cause underlying common cortical sources to project to the scalp in different topographies across subjects (Michel and Murray, 2012). To account for this and other related consequences of EEG volume conduction, we performed EEG source estimation. Using digitized electrode locations and individual head models constructed from subjects’ anatomical MRI data (Thielscher et al., 2015), subject-specific forward models of signal propagation from dipoles distributed over and oriented perpendicular to the cortical surface to electrodes on the scalp were constructed (Gramfort et al., 2010; Kybic et al., 2005). Inverse kernels mapping measured scalp EEG activity to underlying cortical sources were estimated using weighted minimum-norm estimation as implemented in Brainstorm (Tadel et al., 2011). Six cortical regions of interest (ROI) were constructed for each individual, one per stimulation site. Each ROI consisted of all cortical mesh vertices within a radius of 20 mm from the stimulation site (Fig. 1F). Source TEP time series were extracted for each stimulation condition, using the ROI corresponding to the site of stimulation. Trial-averaged TEPs were calculated similarly as for the sensor space data.

#### 2.4.4. Statistical analyses

To test the effect of TMS location and coil orientation, we performed separate two-way repeated-measures analyses of variance (rmANOVAs; factors: stimulation location and coil angle) with Geisser-Greenhouse correction for the EL-TEP (sensor and source space) and artifact amplitudes (Fig. 2C, G). Conditions with two blocks were averaged for this analysis. Normality was evaluated using the Shapiro–Wilk test (QQ plots in Fig. S9). To further explore how stimulation location affected the EL-TEPs and early artifacts, we divided the targets into anterior-posterior (targets t1-4) and the medial-lateral (targets t6, t2 and t5) axes (Fig. 2D, E, H and I). For EL-TEPs and artifacts, we conducted separate *post hoc* paired t-tests (corrected for multiple comparisons) for posterior-to-anterior (i.e. t1 versus t4) and medial-lateral (i.e. t6 versus t5) extremes. Then, to better understand how TMS-evoked artifacts relate to EL-TEPs, targets were categorized into those with low artifact (t1, t2 and t6) and high artifact (t3, t4 and t5) based on a median split of averaged artifact amplitudes. The average EL-TEPs for low versus high artifact targets were compared using a two-tailed paired t-test (Fig. 2J). To test uniqueness of each target’s evoked whole-brain spatio-temporal activations (in sensor space), the difference of each target was statistically compared to all other targets using cluster-based permutation statistics (Maris and Oostenveld, 2007) (cluster threshold<0.05 (two-tailed); two-tailed dependent samples t-test with alpha<0.025; 10000 randomizations; time included: 20-50 ms), as implemented in FieldTrip toolbox (Oostenveld et al., 2011) (Fig. 4D).

**Figure 2.**
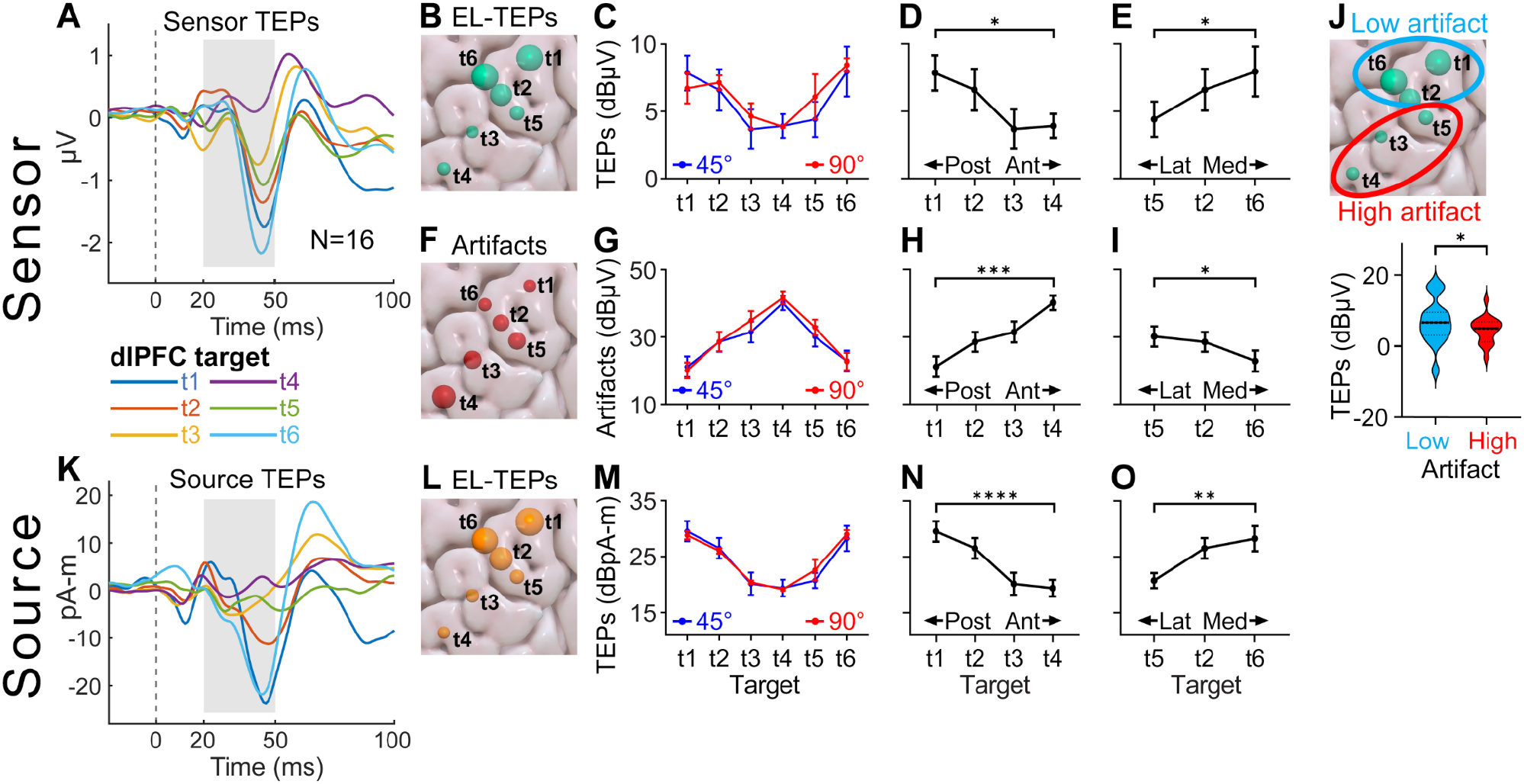
EL-TEPs and artifacts are sensitive to TMS target within the dlPFC. A) Grand average (N=16) EL-TEPs for each stimulation target (ROI: left prefrontal cortex, 45° coil angle). Grey area represents the time window (20 to 50 ms) in which the EL-TEP amplitude is calculated. B-E) Relationship between stimulation targets and EL-TEPs (45° coil angle for B, D and E). In panel B, marker size represents the EL-TEP across subjects. F-I) Same as B-E but for artifacts. J) Relationship between artifact magnitude and EL-TEP amplitude. K) Grand average (N=16) EL-TEPs for each stimulation target in source space (45° coil angle). L-N) EL-TEP amplitudes in source space (45° coil angle for L, N and O). *In C-E, G-I and M-O lines represent mean* ± *SEM (N=16). Asterisks denote the significance level of the post hoc t-tests (corrected for multiple comparisons, N=16): *P<0.05, **P<0.01, ***P<0.001, ****P<0.0001*.

#### 2.4.5. Quantification of noise and SNR

As part of a supplemental analysis, we examined how location within the dlPFC related to the within-block variance of EL-TEPs. We employed the bootstrapped standardized measurement error (bSME) (Luck et al., 2021) to quantify uncertainty in TEP peak-to-peak amplitude with 10000 bootstrap repeats per estimate, accounting for trial-to-trial variation in responses and number of trials aggregated (Fig. S10-11). With bSME calculated on logarithmic (dBμV) amplitude values, we obtained bSME measures in units of dB, and calculated the signal-to-noise ratio (SNR) for each target as the mean response amplitude minus the estimated bSME value for the same condition (Fig. S12).

## 3. Results

### 3.1. EL-TEPs and artifacts are sensitive to TMS target location across the dlPFC

#### 3.1.1. EL-TEPs are sensitive to TMS target

We compared EL-TEP amplitude across targets and angles using two-way rmANOVAs. We found a main effect of target location on EL-TEPs in sensor space (*F*_2.69,40.36_=4.45, *P*=0.011) but not coil angle (*F*_1,15_=1.94, *P*=0.18; Fig. 2C). We observed a significant interaction of stimulation location and coil angle on EL-TEPs (*F*_3.60,54.00_=2.98, *P*=0.031). Target t6 with 90° coil angle produced the largest EL-TEPs on group level. Since the main effect of coil angle was non-significant, for further TEP analyses we focus here on 45° angle conditions (results for the 90° coil angle are shown in Fig. S13). In addition to sensor space, we compared EL-TEP amplitude across targets and angles in source space. We observed a significant main effect of location (*F*_3.33,49.97_=14.74, *P*<0.0001) but not of coil angle (*F*_1,15_=0.49, *P*=0.50) or of interaction of stimulation location and coil angle (*F*_4.46,66.90_=1.76, *P*=0.14; Fig. 2G).

#### 3.1.2. Posteromedial targets produce larger EL-TEPs than anterolateral targets

As a follow-up analysis, we asked whether moving the coil along anterior-posterior and medial-lateral axes would produce different TEP amplitudes. The most posterior (t1) and most anterior (t4) targets had significantly different EL-TEP amplitudes in sensor space (*t*_15_=3.38, *P*=0.017; Fig. 2D) as well as in source space (*t*_15_=6.48, *P*<0.0001; Fig. 2N). The most medial (t6) and most lateral (t5) targets also were significantly different in EL-TEP amplitudes both in sensor (*t*_15_=2.98, *P*=0.028; Fig. 2E) and source space (*t*_15_=3.78, *P*=0.0090; Fig. 2O).

#### 3.1.3. Artifacts are sensitive to TMS location but not coil angle

We next explored the effect of early artifacts across dlPFC targets and coil angles (Fig. 2G). We observed a strong main effect of target location on artifacts (two-way rmANOVA, *F*_3.76,56.34_=27.73, *P*<0.0001), no main effect of coil angle (*F*_1,15_= 0.85, *P*=0.37), and no interaction between location and coil angle (*F*_2.85,42.77_=0.77, *P*=0.51). Since the effect of coil angle was non-significant, for further artifact analyses we here focus on 45° angle conditions (results for the 90° coil angle are shown in Supplementary Material).

#### 3.1.4. Posteromedial targets produce smaller artifacts than anterolateral targets

As a follow-up analysis, we asked whether moving the coil along anterior-posterior and medial-lateral axes produced different amounts of artifact. We found that the most posterior (t1) and most anterior (t4) targets had significantly different artifacts (*t*_15_=5.88, *P*=0.00015; Fig. 2H). The most medial (t6) and most lateral (t5) targets also differed (*t*_15_=2.91, *P*=0.028; Fig. 2I), with the most lateral exhibiting larger artifacts.

#### 3.1.5. Targets with smaller artifacts produce larger TEPs

To further understand the relationship between early artifact and TEP strength, which could inform online TEP optimization, we compared EL-TEP amplitudes (sensor space) between large-artifact and small-artifact targets. After performing a median split (see Methods), targets with small artifacts produced larger EL-TEPs than targets with large artifacts (*t*_15_=2.22, *P*=0.042; Fig. 2J).

#### 3.1.6. Level of noise is not sensitive to stimulation target

As a part of Supplementary analysis, we quantified bSME for each sensor space condition (Fig. S10). We observed no main effect of target location on bSME in μV units (two-way rmANOVA, *F*_3.22,48.26_=1.74, *P*=0.17; Fig S10A), no main effect of coil angle (*F*_1,15_=0.015, *P*=0.91), and no interaction between location and coil angle (*F*_3.11,46.59_=0.37, *P*=0.78). Moreover, we observed no main effect of target location on bSME in dBμV units (two-way rmANOVA, *F*_3.39,50.77_=0.63, *P*=0.62; Fig S10B), a significant main effect of coil angle (*F*_1,15_=5.49, *P*=0.033), and no interaction between location and coil angle (*F*_2.85,42.78_=0.40, *P*=0.74). Since SME was relatively consistent across conditions, and SNR (in dB) was calculated as the difference between TEP amplitude and SME, the group-level SNR had a similar distribution across targets (Fig. S12) compared to the TEP amplitude data (Fig. 2C), i.e. targets with higher TEP amplitudes yielded higher SNR. Across all stimulation targets and subjects, bSME was strongly negatively correlated with EL-TEP amplitude when using dBμV values (*R*=-0.71; Fig. S11A); when using linear EL-TEP values (μV), bSME was positively correlated with EL-TEPs (*R*=0.78; Fig. S11B). Together, these relationships suggest a mix of additive and multiplicative noise in these EL-TEP measurements. Overall, stronger TEPs were associated with higher SNR at the group level.

In summary, within the dlPFC 1) EL-TEPs and artifacts were sensitive to target location and 2) posterior and medial targets produced larger EL-TEPs and smaller artifacts compared to anterior and lateral targets.

### 3.2. Individual variation in EL-TEPs across dlPFC targets

Individual variation of EL-TEPs need to be characterized in order to understand if personalization is needed. To do so, we evaluated the target / angle combination that produced the strongest EL-TEP in each individual (in sensor space). We observed high variability across subjects – only four out of 16 subjects had an individual optimal target / angle combination that matched the group-level best combination (target t6, 90° coil angle; Fig. 3A and B). We compared the individual best EL-TEPs to group best TEPs (t6, 90° coil angle), and TEPs in standard clinical targets (target t3 close to BA9 definition (Rajkowska and Goldman-Rakic, 1995) and target t4 with activity which is most negatively correlated to the sgACC (Fox et al., 2012)). Group best target produced 102% larger EL-TEPs compared to the standard clinical targets. Selecting the target / angle combination that yielded the strongest EL-TEPs for each subject demonstrated a 36% increase over group best and 174% over the EL-TEP from standard clinical targets (in μV values; Fig. 3D).

**Figure 3.**
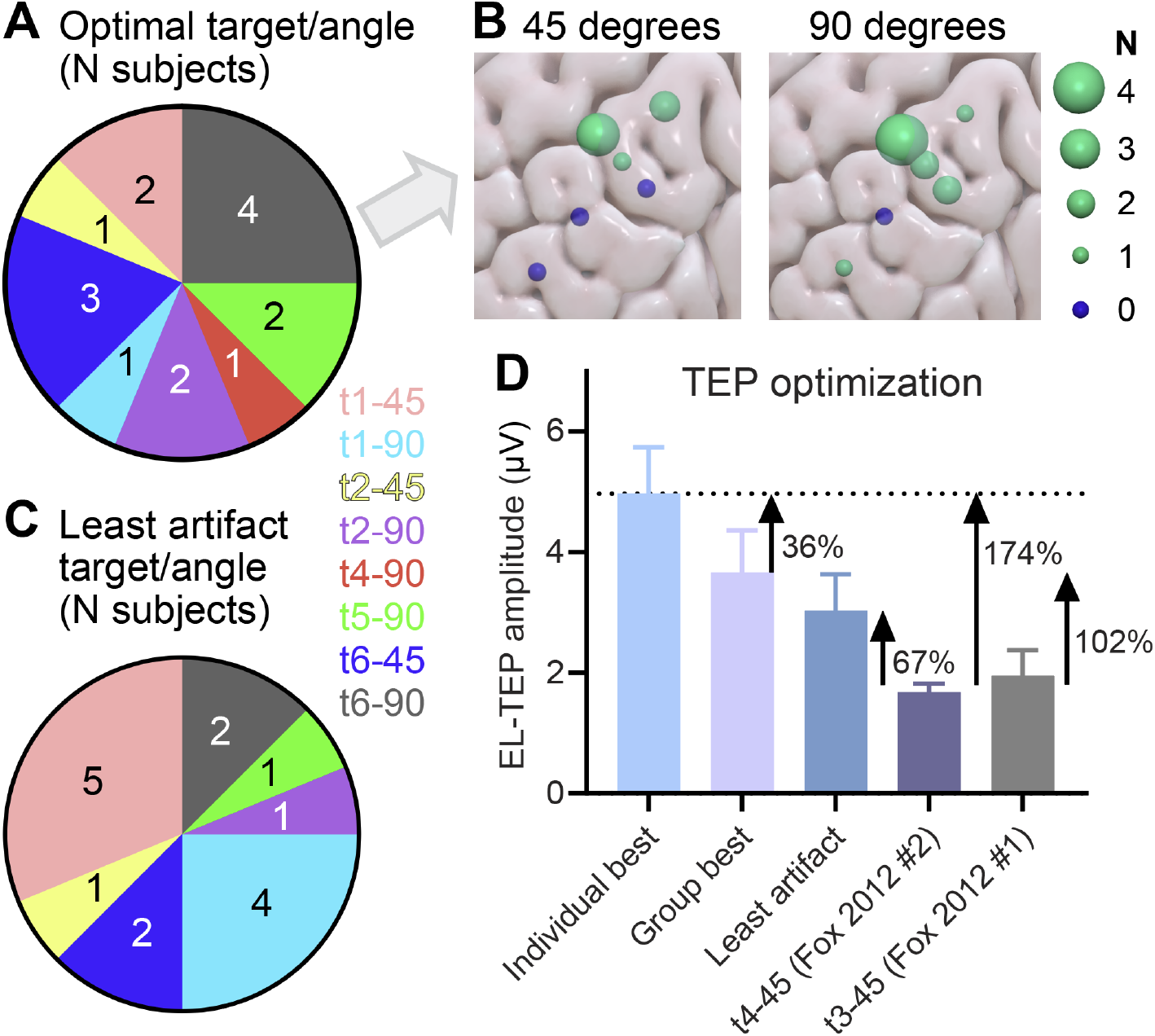
Target selection maximizes the EL-TEP. A) Pie chart of individual best target / angle combinations (the combination that produced largest EL-TEPs in sensor space). Number in each slice represents the number of subjects having a given target / angle combination as their optimal. B) Pie chart of target / angle combinations that produced the smallest artifacts. C) Number of subjects with each target location as their optimal, shown on cortical surface. D) EL-TEP as a function of different targets. ‘Group best’ EL-TEP is larger than the EL-TEP responses to targets negatively correlated to sgACC (t4-45, which has the same coordinates as suggested in Fox et al. 2012) and t3-45 (which is located between ‘BA9 Definition’ of Rajkowska et al. 1995 and another target with activity negatively correlated to sgACC (Fox et al., 2012)). The group best also yields larger EL-TEPs than the ‘least artifact’ targets. *In D bars represent mean ± SEM (N=16)*.

Moreover, we retrospectively explored which target / angle combination produces the smallest artifacts in each subject (Fig. 3C). For four out of 16 subjects, the target / angle combination that produced the smallest artifact matched the combination that produced largest TEPs in that individual. Individually selected least artifact combinations produced 67% larger TEPs over the average of standard clinical targets (in μV values; Fig. 3D). Moreover, the group best target / angle combination (t6, 90° coil angle) produced 21% larger TEPs compared to the smallest artifact combinations. In summary, a group best target / angle for TMS studies will improve EL-TEPs, and a personalization approach (minimizing the artifacts or maximizing TEPs online) may provide added benefit.

### 3.3 Differences in brainwide responses across dlPFC targets

As an exploratory analysis to characterize stimulation target specificity of the brainwide neural response to TMS with fewer assumptions about relevant electrode locations, we compared the sensor-space TEP response at each target (t1-t6) to the average responses to all other targets (45° coil angle for all). Cluster-based permutation testing of the differences between each of these sites and the average responses at all other sites between 20-50 ms revealed significant differences (Fig 4B-C). Significant target-specific activations were found for each target except t5, suggesting that brainwide spatiotemporal patterns in the 20-50 ms response window are sensitive to TMS target in the dlPFC.

**Figure 4.**
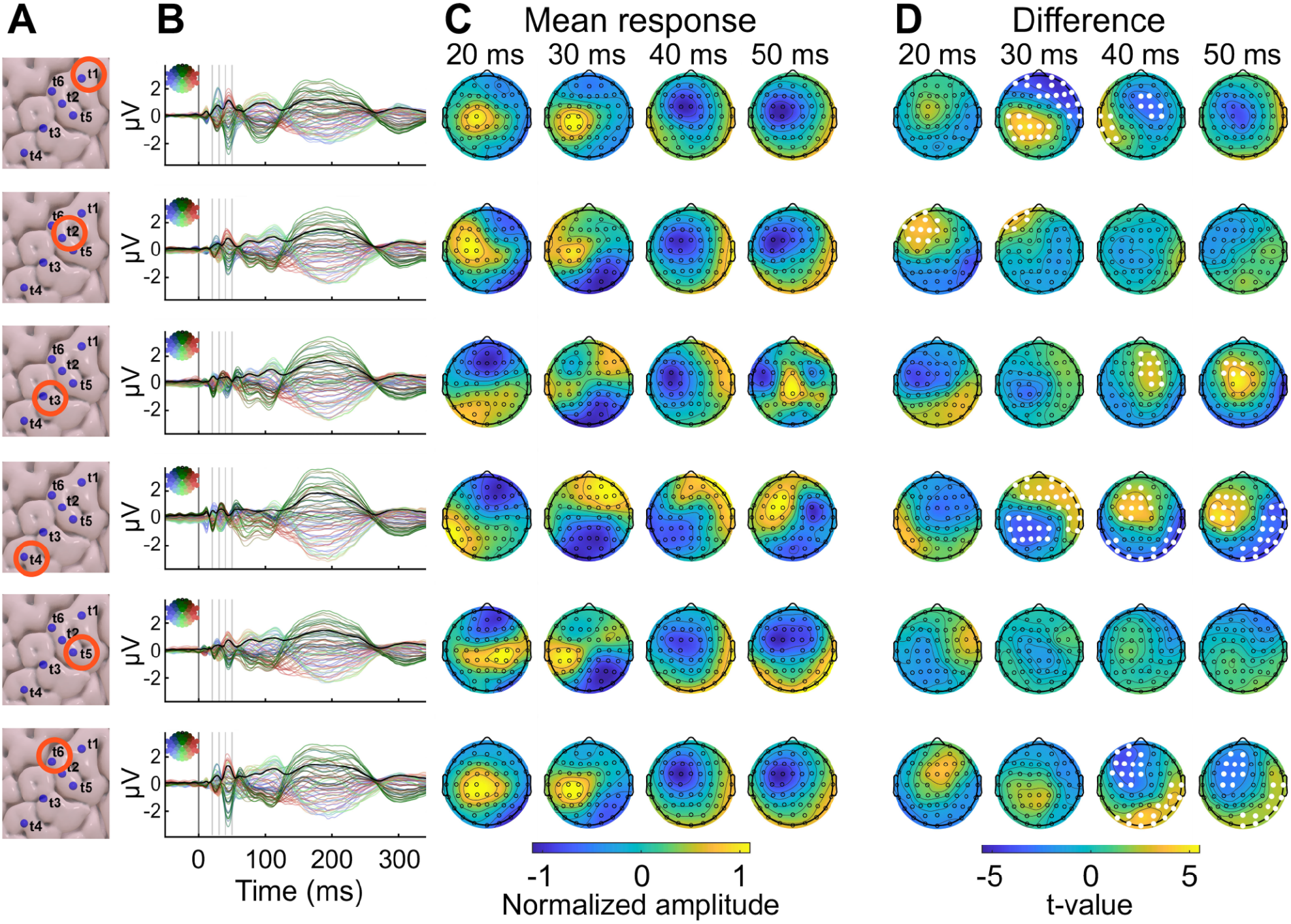
Stimulation of dlPFC targets elicit unique whole-brain spatio-temporal activation patterns in sensor space. A-C) Group (N=16) location (A), butterfly plot (B), and scalp topographies (C) for all targets (all at 45° angle). D) Spatio-temporal differences between a selected target and all other targets, as implemented by cluster-based permutation testing. Electrodes belonging to statistically significant clusters are highlighted as white dots (*P<0.001*).

## 4. Discussion

### Summary of findings

In the current study we sought to better understand how the EL-TEP, a causal measure of cortical excitability, varies across TMS coil target locations and coil angles when stimulating the dlPFC. Our findings were as follows: 1) EL-TEPs and artifacts varied significantly as a function of dlPFC target. Specifically, lateral and anterior targets produced smaller EL-TEPs and larger artifacts, whereas medial and posterior targets produced larger EL-TEPs and smaller artifacts. The group best target yielded 102% larger EL-TEP compared to standard targets.; 2) Individual offline optimization of target and angle enhanced the EL-TEP by 36%; 3) All targets except the most lateral target (t5) showed different whole-brain spatio-temporal responses in the EL-TEP time window compared to the other targets; 4) EL-TEPs contained a mix of additive and multiplicative noise, and stronger TEPs were associated with higher SNR at the group level. In summary, the study furthers our understanding of this causal metric of cortical excitability across the dlPFC. This work suggests that effects of TMS treatment could be confidently monitored with EL-TEPs in the posterior-medial aspects of the dlPFC and that a personalized, real-time approach to targeting may further improve the signal quality of the EL-TEP.

### EL-TEP as a function of dlPFC target and coil angle

To date, little is known about how noninvasive brain measurements, including the EL-TEP, vary across stimulation targets and coil angles within the dlPFC. This information is critical to effectively monitor the neural effects of TMS treatment. Here, we observed a large variability in EL-TEP strength across different targets and coil orientations. Specifically, EL-TEPs were up to two times larger in posterior-medial regions compared to anterior-lateral regions, and in regions where EL-TEPs were larger, early artifacts were smaller (Fig. 2). These results may be due to the fact that the large frontalis muscle is typically located in closer proximity with anterior targets, whereas the temporalis muscle is located close to the lateral targets. It is worth noting though that there is large inter-subject variability in the anatomy of frontalis muscle (Costin et al., 2015), and on an individual level the relationship between EL-TEPs and muscle artifact was only moderately inversely correlated (Fig. S14). Whether this difference in EL-TEP variability across targets represents differences in true cortical excitability or strength differences from the muscle artifact masking EL-TEPs is unclear. Further work is needed to explore how coil angle should be modified based on target location to optimize EL-TEPs. Future studies should also focus on relating differences in EL-TEPs across targets and angles to differences in anatomical structure (e.g. gyral folding) and downstream functional networks to enhance its biological validity. EL-TEPs have been linked to GABAergic inhibition, and related to both depression (Voineskos et al., 2019) and treatment response to TMS (Eshel et al., 2020; Voineskos et al., 2021). Multiple lines of evidence also suggest that EL-TEPs have intracranial neural correlates (Huang et al., 2019; Keller et al., 2018). However, more work is needed to better understand the relationship between the strength of EL-TEPs to downstream functional brain networks. Before testing clinical utility of EL-TEPs, the optimized TEPs need to be evaluated for reliability across multiple visits and sensitivity to the neural effects of TMS, most importantly in patients. In summary, this work sheds light on the dlPFC subregions that can be confidently monitored with large EL-TEPs and emphasizes the need for future studies on the biological validity of EL-TEPs.

### Optimization of EL-TEPs

This work suggests promise for a personalized approach for improving the EL-TEP signal. Real-time EL-TEP optimization is an important next step for ensuring high quality EL-TEPs during experiments or treatment. Using a retrospective group-best target / angle combination (target t6, 90° coil angle), EL-TEPs were 102% larger than standard clinical targets (Fig 3C). In comparison, EL-TEPs were enhanced 67% when retrospectively selecting targets / angles that minimize artifacts or by 174% when selecting targets / angles that maximize EL-TEPs (Fig 3C). It is worth considering the feasibility of these three approaches for future experiments. The group approach requires no real-time optimization and yields a strong boost in signal. While the *individualized EL-TEP* approach yields stronger gains, it is logistically much more difficult to execute, as it requires real-time optimization of the EL-TEP. A compromise may be the *individual least artifact* approach. Estimating artifact magnitude typically requires less data and less advanced real-time processing compared to EL-TEP estimation. In our retrospective analysis, the individual least artifact approach provided gains over the standard clinical targets, but was not more effective than the group approach. This is either because 1) the relationship between artifact and EL-TEP is complex (Fig 2) or 2) because a higher resolution in target / angle search would provide significant gains. Furthermore, in the real-time approach, more dimensions can be tested including intensity. This online approach to optimize artifacts and EL-TEPs has been shown to be feasible, at least with a custom-made TMS setup and stimulating pre-supplementary motor region and sufficient time for experimentation (Tervo et al., 2021). Online optimization was not utilized in this experiment as in this study we first wanted to characterize the EL-TEPs and artifacts at fixed locations and angles. Future work will include building out this online optimization approach and prospectively testing its benefit to measure robust and reliable EL-TEPs in prefrontal regions. An important consideration in future studies will be the balance between the extra experimental time needed to optimize these EL-TEPs and the strength of improvement. If the optimization protocol produces only a modest boost, for some studies utilizing the group-best (target t6, 90° coil angle) target may yield strong enough results. For others, only short online optimization with granular adjustments in target, angle, and intensity may be worthwhile.

### Limitations and future directions

Several limitations are worth discussing. First, our choice of brain targets was driven by equidistant sampling of the middle frontal gyrus as well as the desire to probe the anterior-posterior and medial-lateral directions. Future work should follow up on these findings and include analysis of the inter-target variability of the EL-TEP 1) at finer spatial resolution and 2) at additional anterior-medial targets. A second limitation is that we only used two coil angles. Future work should explore how other coil angles affect the EL-TEP. Developing online algorithms to fully automate target / angle search space within and outside of the dlPFC could prove valuable but is outside the scope of this work. Third, it is worth considering the influence of our E-field modeling approach on the results, as offline E-field modeling has its own limitations. Although our results suggest that absolute intensity differences did not contribute significantly to EL-TEP strength (Fig. S7), E-field modeling could influence the effective intensities used to probe different targets (Drakaki et al., 2022; Saturnino et al., 2021). In the future as these models continue to improve, it is worth re-examining these targets with more sophisticated E-field matching.

The EL-TEP time window was *a priori* chosen based on previous reports of neurophysiological effects of TMS in this time window (Esser et al., 2006) and potential clinical relevance (Eshel et al., 2020; Voineskos et al., 2021, 2019). In addition, this time window was chosen to avoid off-target sensory components that are usually reported between 100 and 200 ms (Ross et al., 2022b). It is worth noting that there are potential sensory responses even in this 20-50 ms time window (A novel EEG paradigm to simultaneously and rapidly assess the functioning of auditory and visual pathways - PubMed, n.d.; Davis and Zerlin, 1966; Davis, 1939; Geisler et al., 1958; Knight et al., 1980; Näätänen and Picton, 1987; Vanni et al., 2004), as auditory stimuli can elicit neural responses in EEG within 10 to 60 ms after stimulus onset (for a review, see (Kappenman and Luck, 2011)). However, the EL-TEP has been shown to be unaffected by sensory suppression techniques, supporting the notion that sensory contributions during this time period is likely minimal (Ross et al., 2022a, 2022b). Further, we implemented auditory suppression techniques (Ross et al., 2022b) and our GMFA time series (Fig. S15) support that condition-specific differences in the 100-200 ms window were not accompanied by similar patterns in the EL-TEP time window. For all these reasons, we feel that it is unlikely that the EL-TEP reported on here is sensory in nature although this possibility cannot be definitively ruled out.

Finally, it is worth considering the implications of the study with respect to TMS treatment. High SNR of the EL-TEP says little about the optimal treatment targets within the dlPFC. Multiple dlPFC treatment targets for depression have been proposed based on symptom profiles (Fried and Nesse, 2015) and network activation (Fox et al., 2012; Weigand et al., 2018), and the objective of the current study was simply to determine which of these targets could be sampled with EL-TEPs at high signal quality. Future work is needed to explore the relationship between the optimized EL-TEP target / angle and optimized TMS treatment.

### Conclusions

This study characterized the EL-TEP across brain targets and angles within the dlPFC. We observed clear group-level trends, including posterior and medial targets resulting in larger EL-TEPs and smaller artifacts compared to anterior and lateral targets. The best group-level target resulted in 102% larger EL-TEPs compared to other standard targets. Consistent with previous reports, we observed high variability across subjects, and our data suggests that this variability can be used to personalize and optimize the EL-TEP on an individual subject basis, leading to an additional >30% boost in signal. In summary, this study furthers our understanding of cortical excitability across the dlPFC. This work suggests that effects of TMS treatment could be confidently monitored with EL-TEPs in the posterior-medial aspects of the dlPFC and a personalized approach to this monitoring may further improve its signal strength.

## Acknowledgements

We extend our deep gratitude to all of our research participants. We would also like to acknowledge the generous contributions of the members of the Personalized Neurotherapeutics Laboratory for helpful feedback on the manuscript and throughout the course of the study. This research was supported by the National Institute of Mental Health under award number R01MH126639, R01MH129018, and a Burroughs Wellcome Fund Career Award for Medical Scientists (CJK). JG was supported by personal grants from the Finnish Medical Foundation, Orion Research Foundation and Emil Aaltonen Foundation. JMR was supported by the Department of Veterans Affairs Office of Academic Affiliations Advanced Fellowship Program in Mental Illness Research and Treatment, the Medical Research Service of the Veterans Affairs Palo Alto Health Care System and the Department of Veterans Affairs Sierra-Pacific Data Science Fellowship.

## Supplementary Material

### Analyses of TEPs and artifacts for 90° coil angle

In addition to the analyses for 45° coil angle (see Results), we also conducted similar follow-up analyses for 90° data. We asked whether using the most anterior targets can lead to significantly different EL-TEP amplitudes than when using the most posterior target. We also asked whether using the most lateral target would lead to different TEP amplitudes compared to the most medial target. The most posterior (t1) and most anterior (t4) targets did not have significantly different EL-TEP amplitudes in sensor space (*t*_15_=0.85, *P*=0.65; Fig. S13A), whereas in source space, there was a significant difference between these two targets (*t*_15_=4.10, *P*=0.0047). The most medial (t6) and most lateral (t5) targets did not have significantly different EL-TEP amplitudes neither in sensor (*t*_15_=0.85, *P*=0.65; Fig. S13B) or source space (*t*_15_=2.44, *P*=0.080).

Similarly as for 45° coil angle, we asked whether moving the coil along anterior-posterior and medial-lateral axes would produce differing amounts of artifact with 90° coil angle. We found that the most posterior (t1) and most anterior (t4) targets had significantly different artifacts (*t*_15_=10.84, *P*<0.000001; Fig. S13C). The most medial (t6) and most lateral (t5) targets also differed (*t*_15_=4.59, *P*=0.0011; Fig. S13D), with the most lateral having larger artifacts.

### Figures and plotting

MATLAB R2021b (Mathworks, USA) and GraphPad Prism 9 (GraphPad Software, USA) were used for plotting. Stimulation targets were visualized using Surf Ice (https://www.nitrc.org/projects/surfice/). For the butterfly plots and scalp topographies (Fig. 4B and C), the data of each stimulation target was averaged across subjects. Global mean field amplitude (GMFA) was also overlaid on each butterfly plot. For each topographic visualization, the spatial distributions were normalized. All figures were assembled using Adobe Illustrator 2022 (Adobe Inc., USA).

**Supplementary Table 1.** Distances between the stimulation sites (MNI space).

| **Distances** | **Point1** | **Point2** |  |
| --- | --- | --- | --- |
| **14.85** | t2 | t1 | a-p |
| **14.85** | t3 | t2 |  |
| **14.85** | t4 | t3 |  |
| **29.71** | t4 | t2 |  |
| **44.56** | t4 | t1 |  |
| **15.36** | t5 | t6 | lat-med |
| **17.95** | t1 | t5 |  |
| **15.39** | t1 | t6 |  |
| **7.68** | t2 | t5 |  |
| **7.68** | t2 | t6 |  |

**Supplementary Table 2.** Stimulation intensities. The table shows rMT values and stimulation intensities (in %MSO) for all subjects and conditions.

| Sub | rMT | 45 degs |  |  |  |  |  | 90 degs |  |  |  |  |  |
| --- | --- | --- | --- | --- | --- | --- | --- | --- | --- | --- | --- | --- | --- |
|  |  | t1 | t2 | t3 | t4 | t5 | t6 | t1 | t2 | t3 | t4 | t5 | t6 |
| 1 | 52 | 64 | 50 | 45 | 51 | 46 | 59 | 64 | 66 | 54 | 58 | 51 | 69 |
| 2 | 47 | 52 | 55 | 50 | 53 | 53 | 57 | 48 | 51 | 56 | 57 | 49 | 53 |
| 3 | 64 | 74 | 87 | 79 | 80 | 84 | 95 | 80 | 88 | 84 | 82 | 89 | 90 |
| 4 | 75 | 83 | 89 | 87 | 83 | 88 | 100 | 78 | 82 | 92 | 80 | 88 | 84 |
| 5 | 70 | 64 | 69 | 63 | 67 | 66 | 76 | 66 | 70 | 67 | 70 | 70 | 73 |
| 6 | 67 | 73 | 68 | 67 | 65 | 69 | 75 | 79 | 76 | 75 | 77 | 72 | 80 |
| 7 | 78 | 88 | 87 | 88 | 100 | 87 | 91 | 84 | 79 | 87 | 84 | 80 | 78 |
| 8 | 67 | 67 | 78 | 70 | 76 | 68 | 80 | 69 | 76 | 73 | 80 | 68 | 76 |
| 9 | 74 | 96 | 97 | 94 | 96 | 91 | 99 | 94 | 98 | 94 | 100 | 96 | 94 |
| 10 | 60 | 57 | 57 | 59 | 63 | 56 | 58 | 59 | 57 | 64 | 65 | 58 | 58 |
| 11 | 74 | 69 | 67 | 71 | 78 | 66 | 68 | 62 | 69 | 79 | 87 | 70 | 69 |
| 12 | 63 | 69 | 68 | 69 | 72 | 67 | 74 | 70 | 78 | 69 | 70 | 74 | 82 |
| 13 | 74 | 88 | 88 | 91 | 97 | 81 | 94 | 87 | 100 | 88 | 91 | 86 | 99 |
| 14 | 74 | 76 | 86 | 86 | 87 | 84 | 95 | 78 | 85 | 90 | 85 | 85 | 91 |
| 15 | 83 | 79 | 93 | 100 | 88 | 82 | 95 | 72 | 82 | 99 | 92 | 77 | 83 |
| 16 | 100 | 87 | 93 | 89 | 90 | 91 | 93 | 79 | 89 | 91 | 89 | 94 | 89 |

**Supplementary Figure 1.**
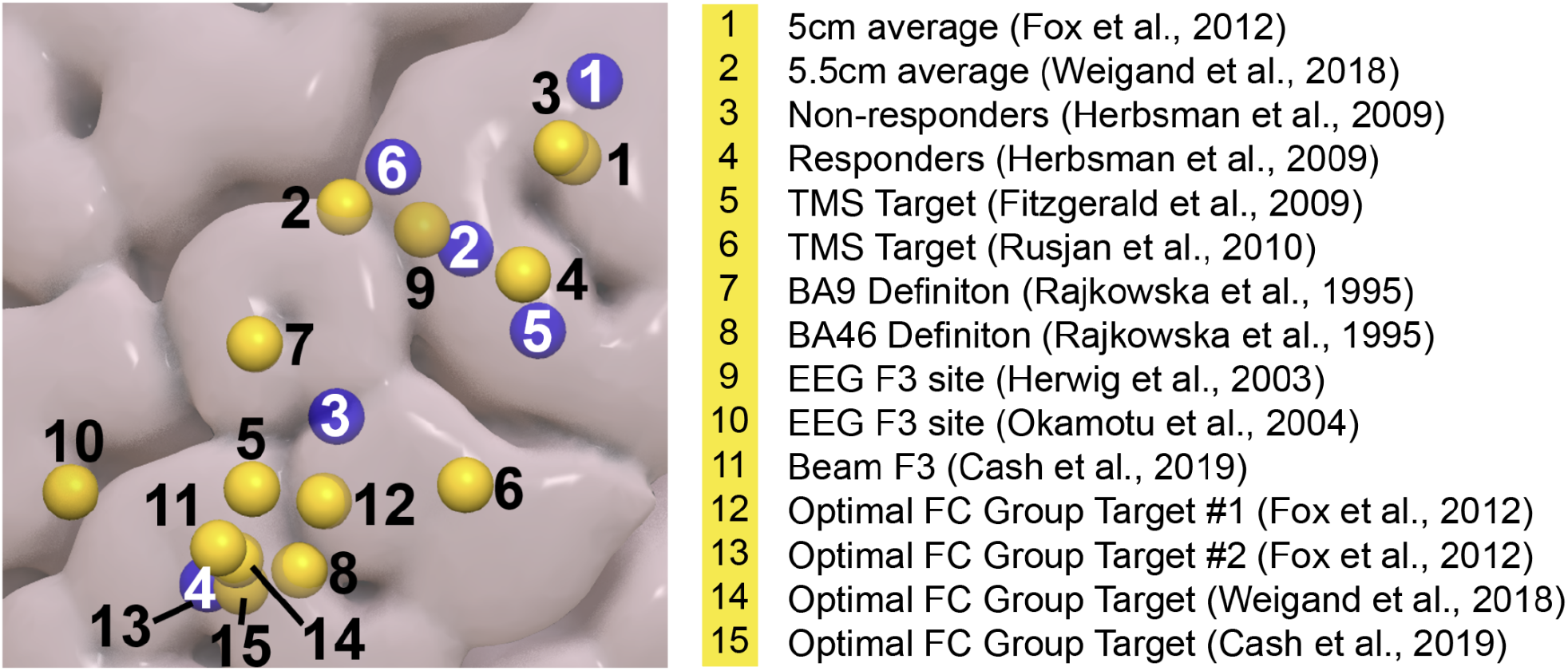
Spatial relationship of the targets used in the current paper compared to earlier depression treatment targets (as summarized by Cash et al., 2021). The treatment target with activity that is most negatively correlated to the sgACC according to Fox et al., 2012 (13 in the figure) has the same MNI coordinates as t4 in the current study. *Blue blobs depict the targets (t1-6) used in the current study, yellow blobs correspond to TMS depression treatment targets*.

**Supplementary Figure 2.**
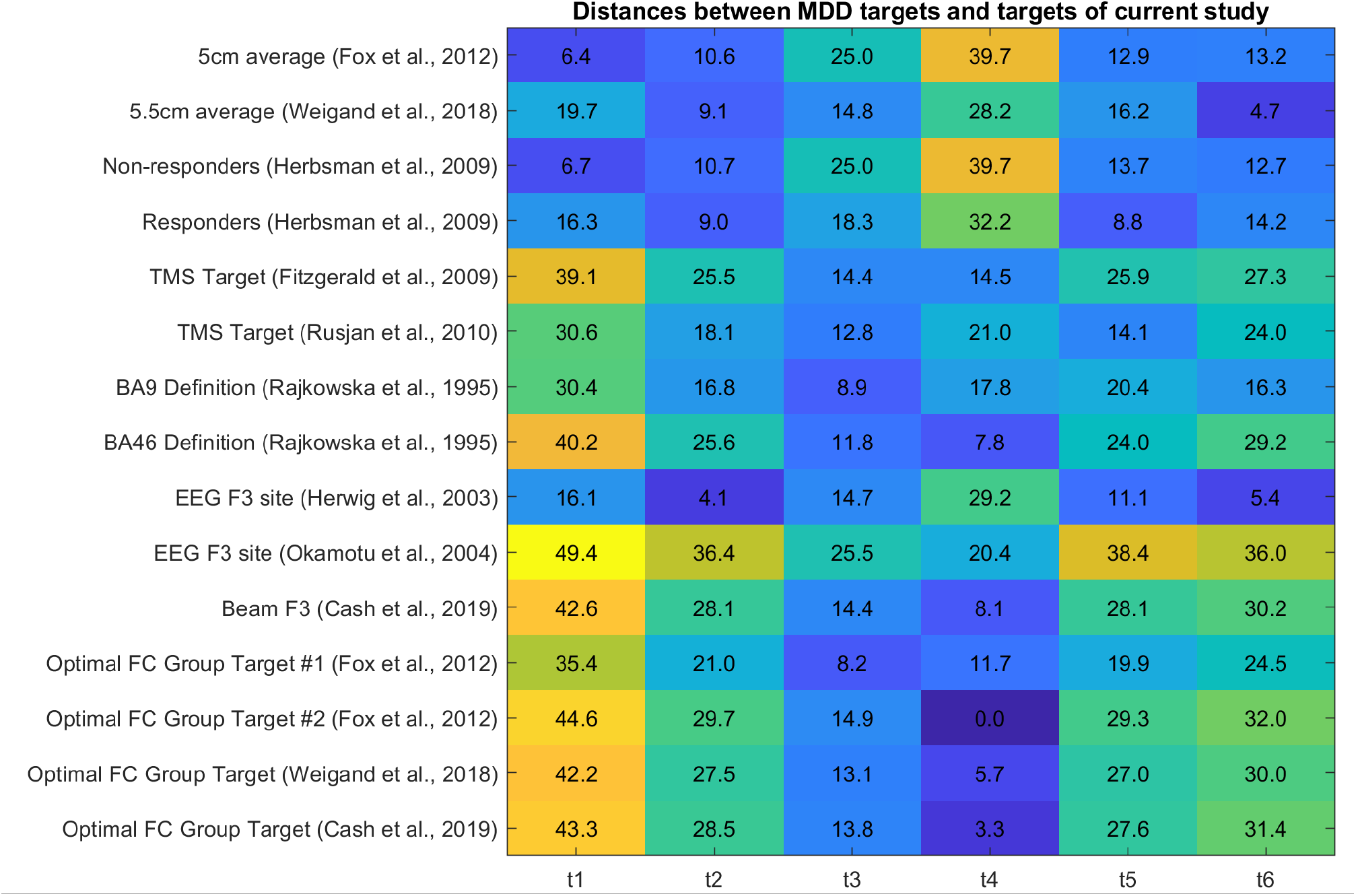
Euclidean distances between the clinical targets (as summarized by Cash et al., 2021) and the targets of the current study.

**Supplementary Figure 3.**
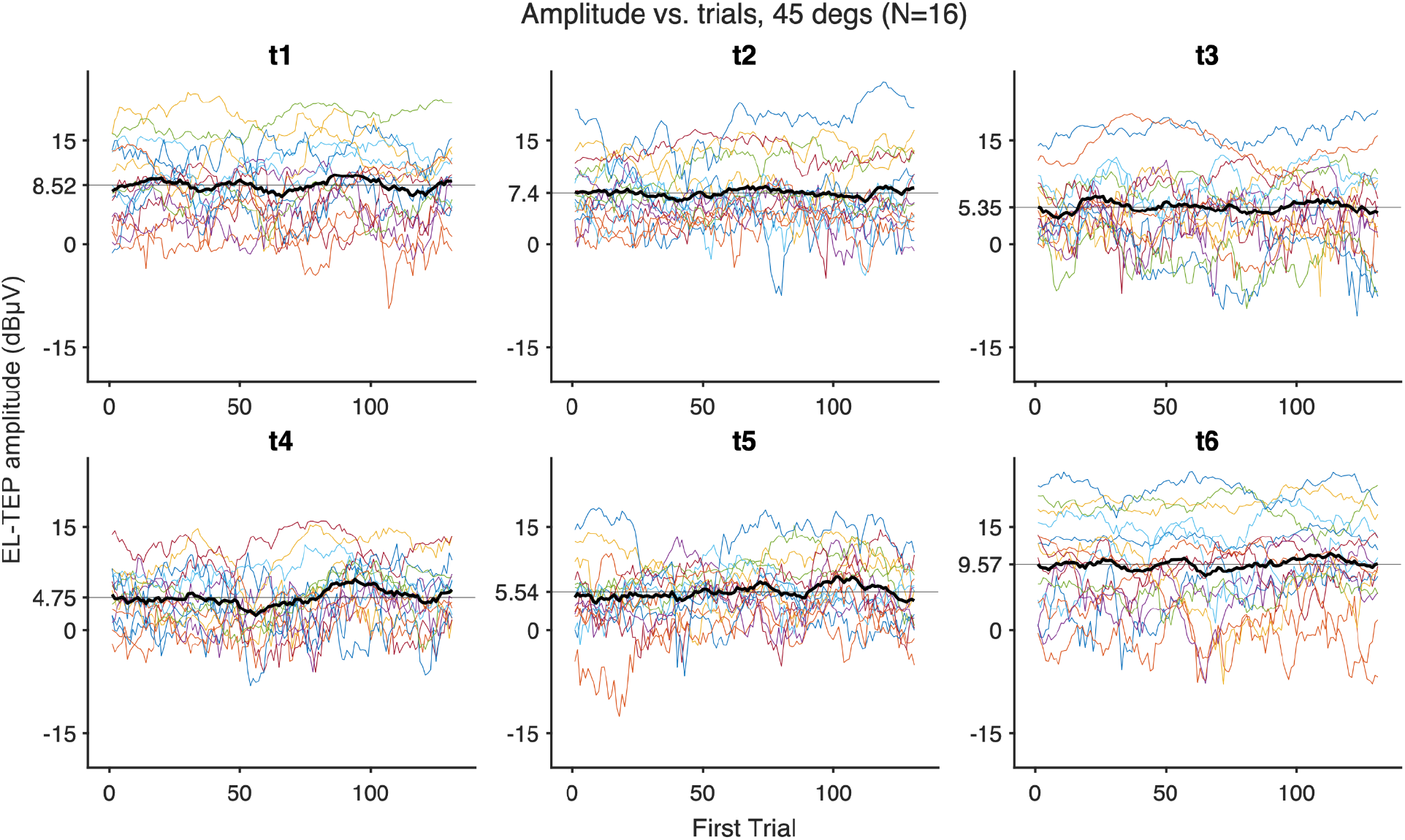
No TEP changes within blocks. Colored lines represent a moving average of TEP amplitude (sensor space), using a sliding set of 20 trials. Black lines show the average across subjects. Vertical straight lines show the average of all moving average TEP amplitudes.

**Supplementary Figure 4.**
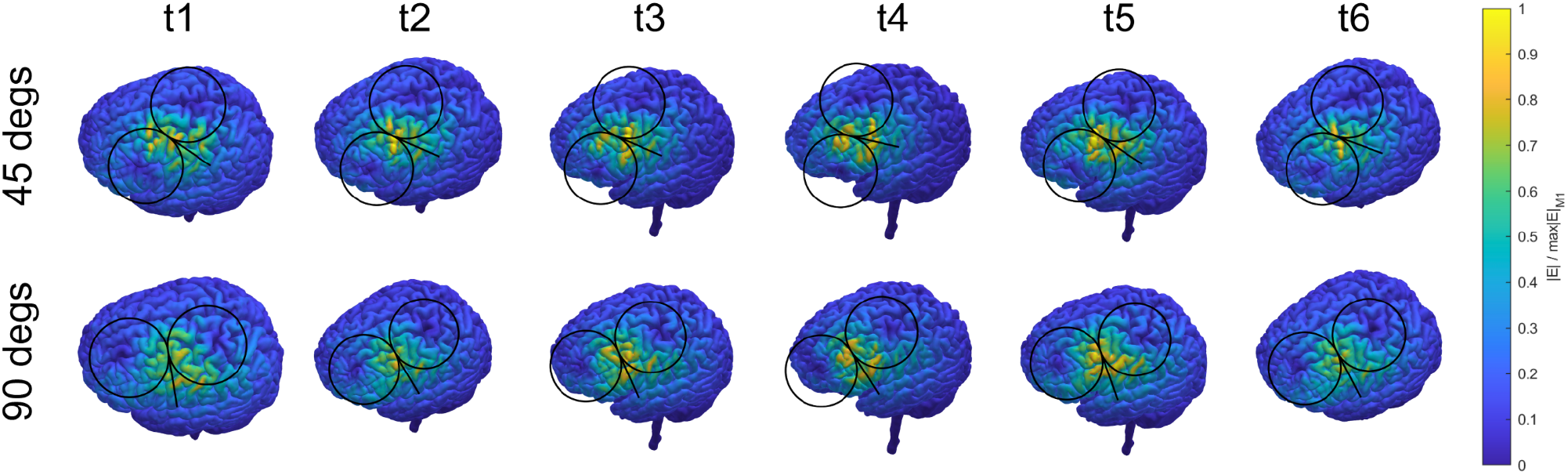
E-fields normalized relative to M1 for a fixed stimulation intensity in a representative subject.

**Supplementary Figure 5.**
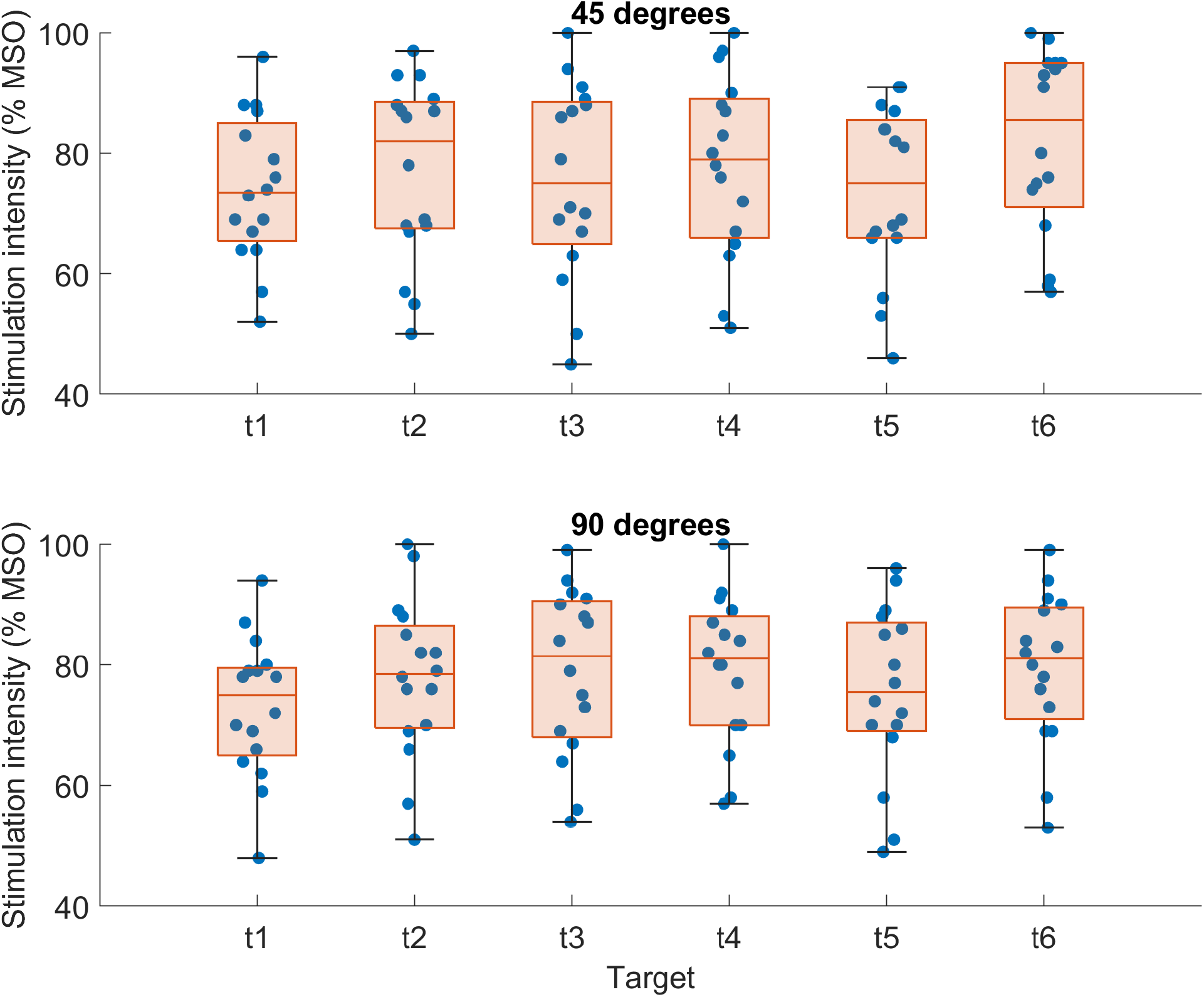
Stimulation intensities in %MSO.

**Supplementary Figure 6.**
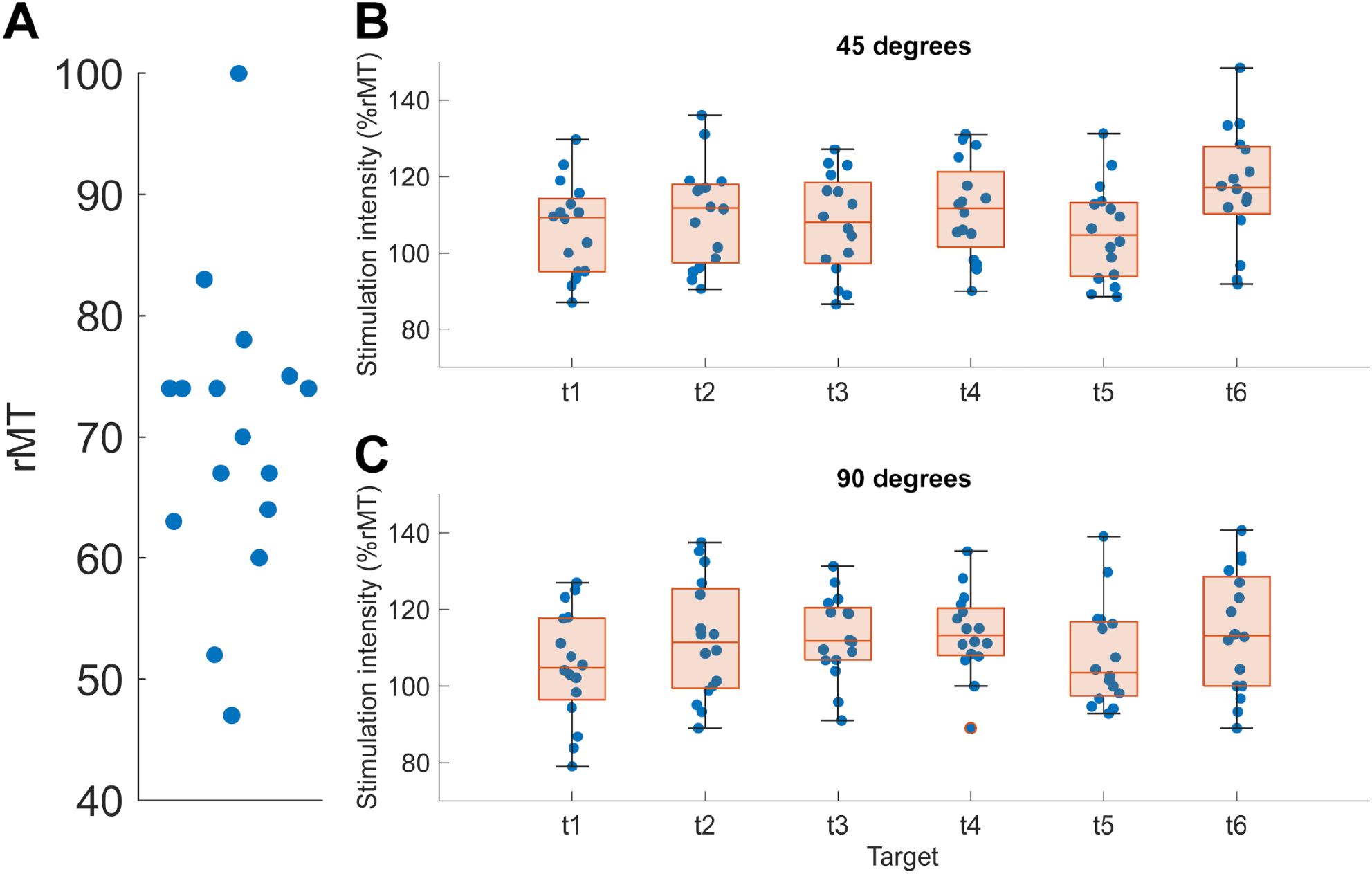
Stimulation intensities. A) Resting motor thresholds (rMTs). B-C) Stimulation intensities as %rMT.

**Supplementary Figure 7.**
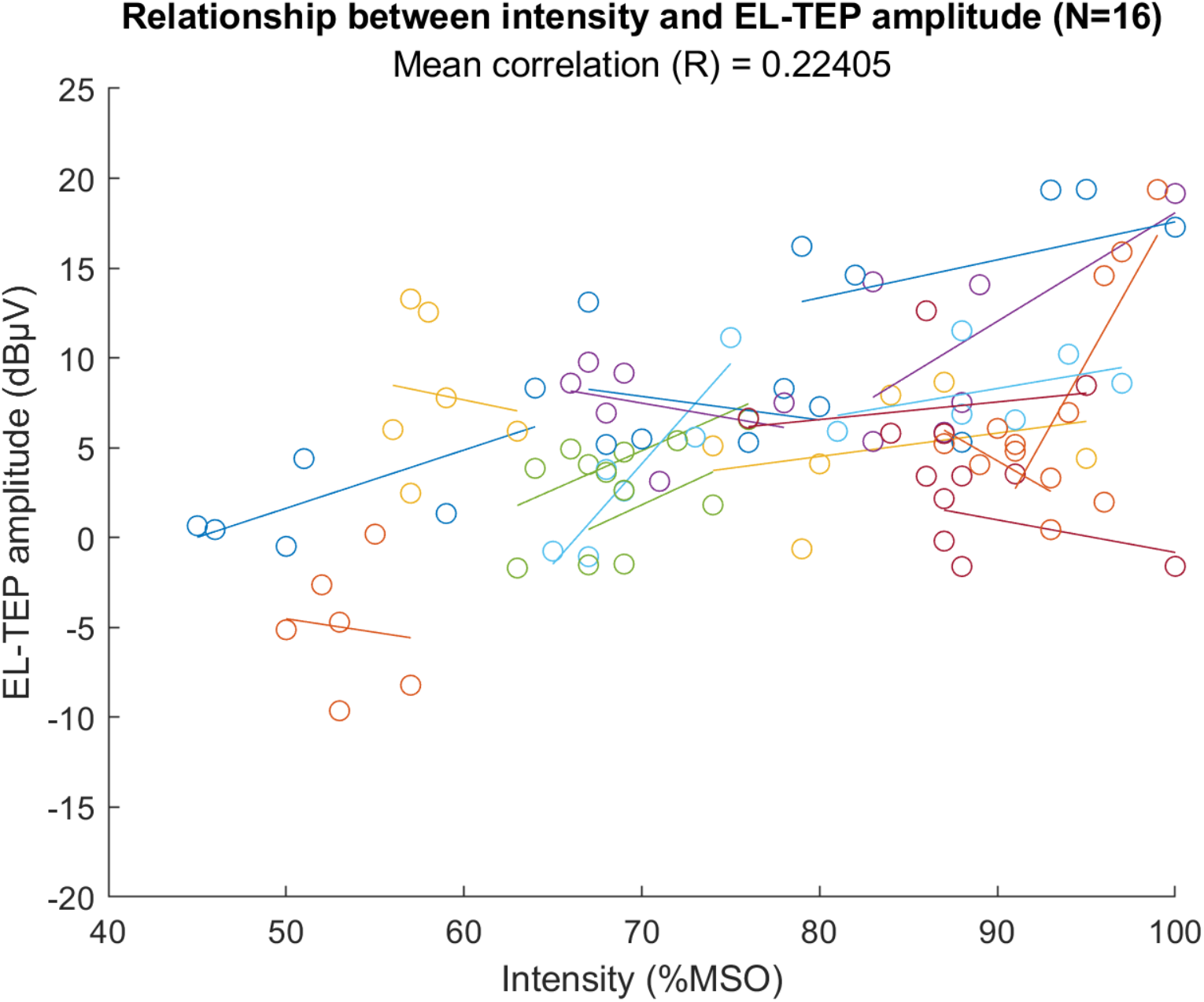
Relationship between stimulation intensity and EL-TEP amplitudes (sensor space). Stimulation intensities were corrected across targets based on calculated maximal induced E-field. Each color represents one subject.

**Supplementary Figure 8.**
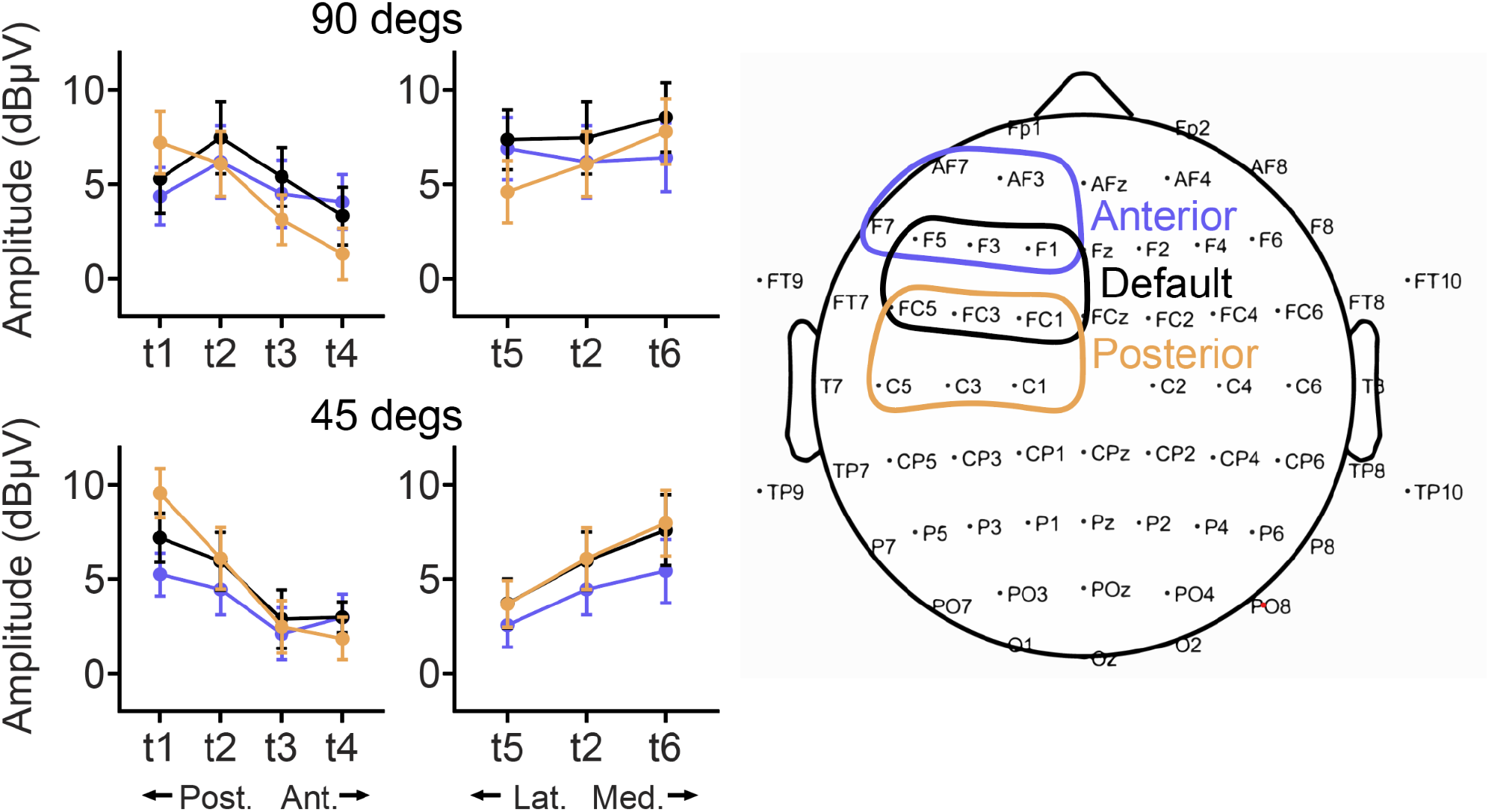
EL-TEP amplitudes for shifted ROIs.

**Supplementary Figure 9.**
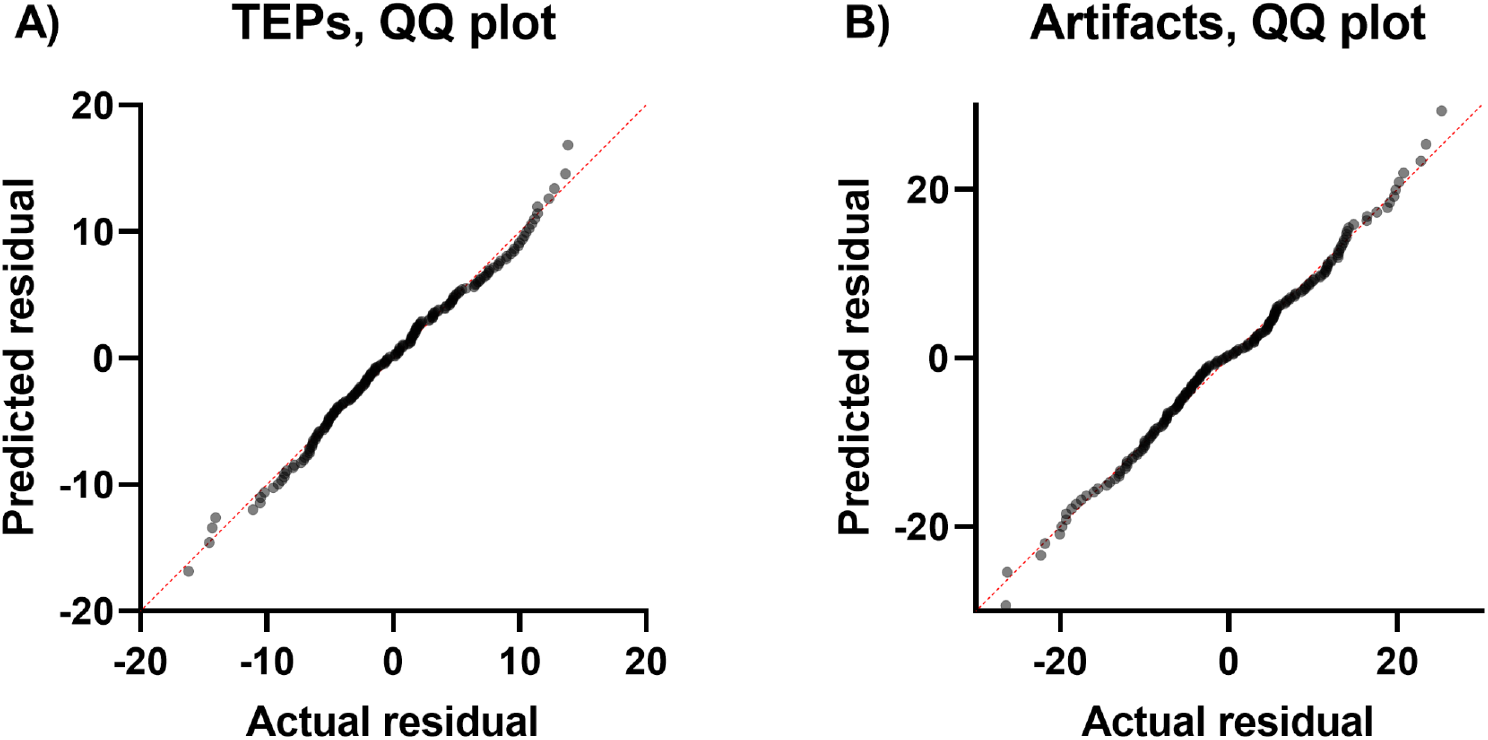
QQ plots for A) EL-TEP (sensor space), and B) artifact distributions.

**Supplementary Figure 10.**
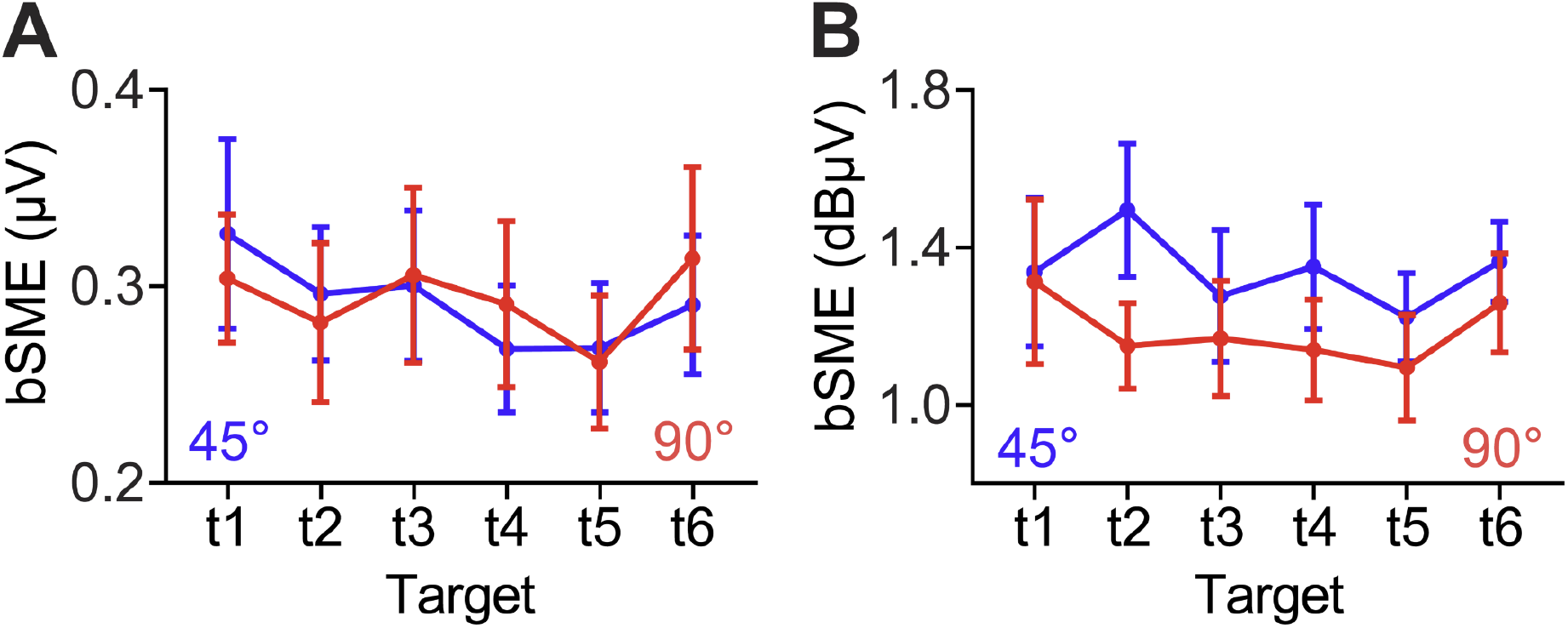
Bootstrapped standardized measurement error (bSME) for each target. We employed the bSME (Luck et al., 2021) to quantify uncertainty in EL-TEP amplitude with 10000 bootstrap repeats per estimate, accounting for trial-to-trial variation in responses and number of trials aggregated. A) bSME for EL-TEPs in μV units. B) bSME for EL-TEPs in dBμV units.

**Supplementary Figure 11.**
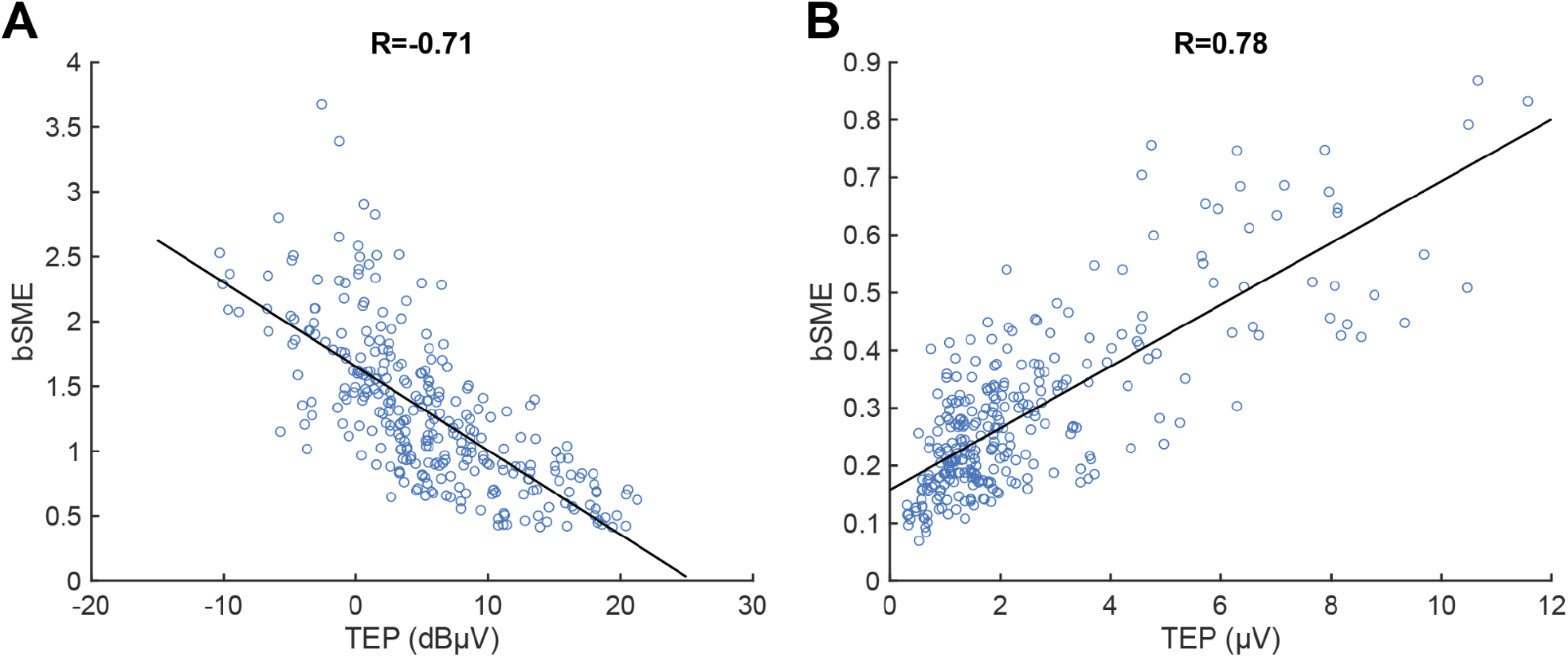
Relationship of bootstrapped standardized measurement error (bSME) and EL-TEP amplitude. Each dot represents an EL-TEP amplitude (sensor space) of one stimulation block (N=16). A) Relationship between bSME and EL-TEPs (in dBμV). B) Relationship between bSME and EL-TEPs (in μV).

**Supplementary Figure 12.**
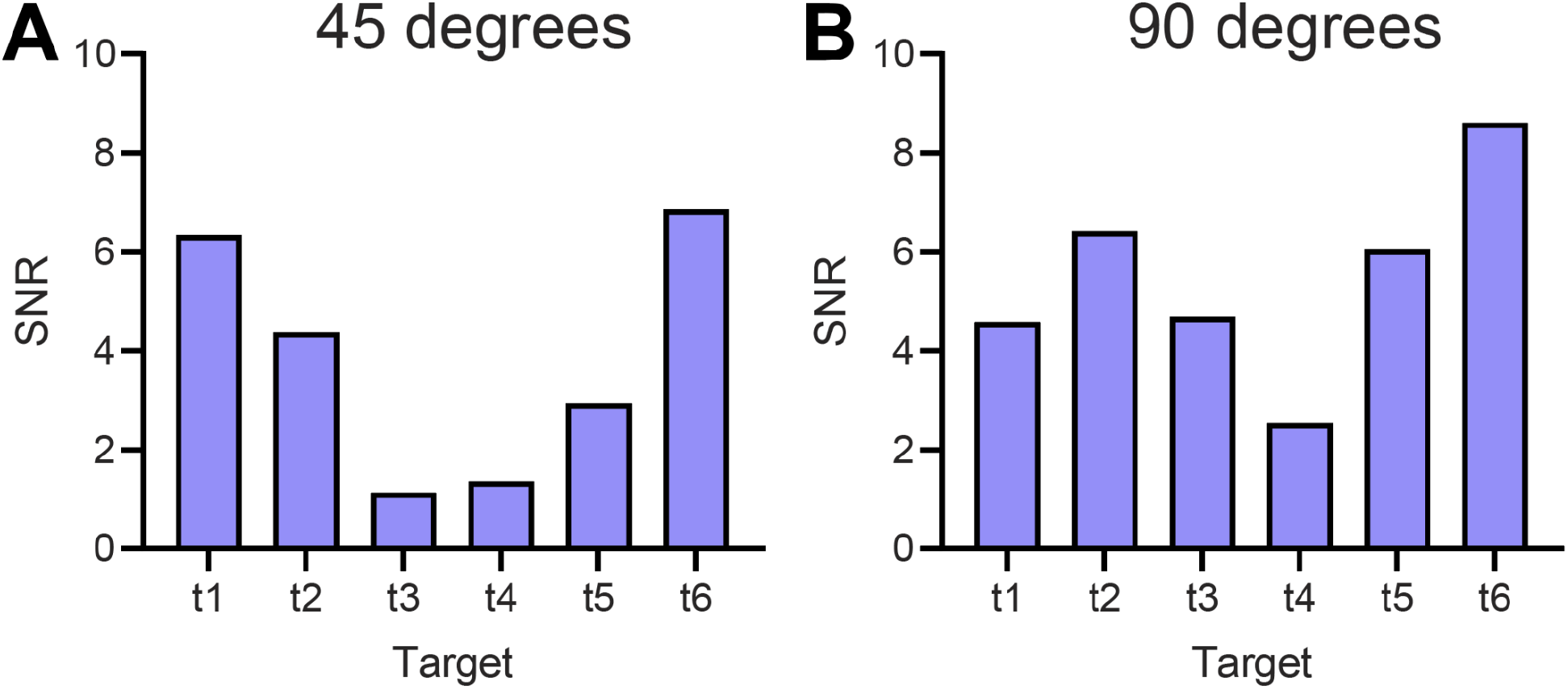
Group-level signal-to-noise ratio (SNR) for each target. SNR was calculated as mean TEP amplitude minus SME_group_, using logarithmic (dBμV) EL-TEP amplitudes in sensor space.

**Supplementary Figure 13.**
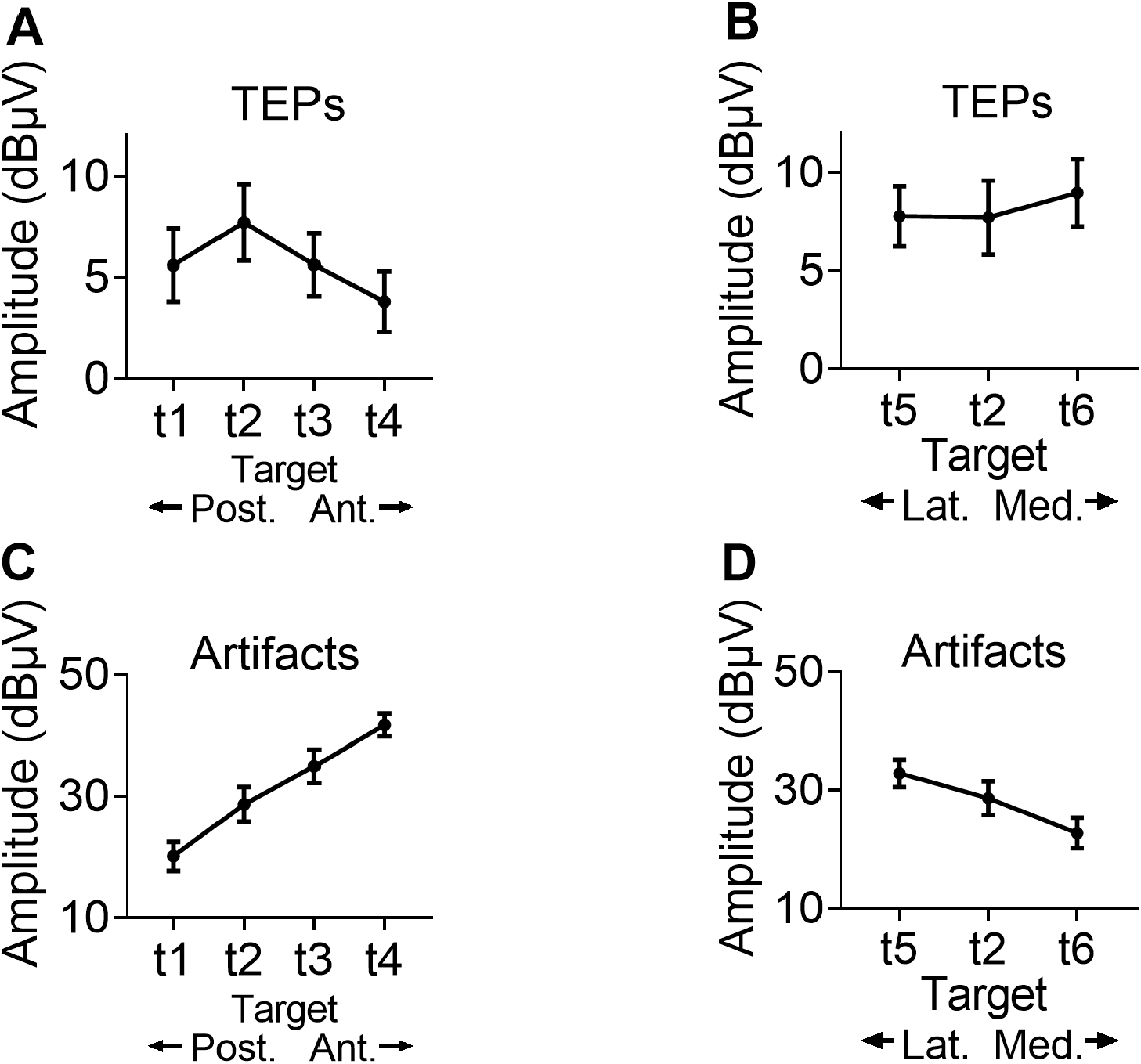
Target-related sensitivity of TEPs and artifacts for 90 degrees coil angle (sensor space). A-B) Relationship between target (t1-t6, x-axis) and EL-TEP amplitude with 90 degrees coil angle. C-D) Relationship between stimulation target and muscle artifact with 90 degrees coil angle.

**Supplementary Figure 14.**
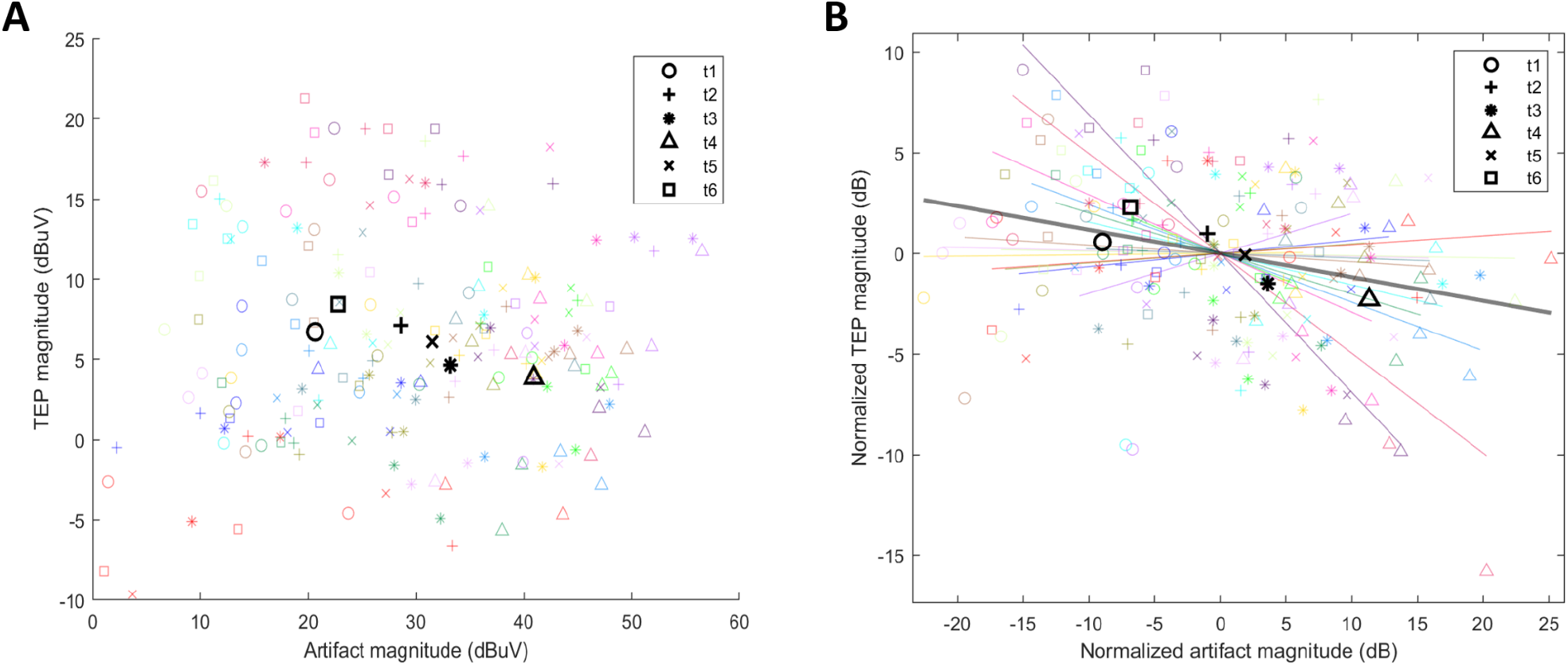
Relationship between EL-TEP amplitudes (sensor space) and artifact amplitudes. Shape of scatter points indicate target site, with one color per subject; black points represent the mean for each target across subjects. Normalized values in B were obtained by subtracting each subject’s mean response magnitude (in dB). Colored lines in B represent linear fit of points across targets within each subject; bold gray line indicates linear fit of the mean values.

**Supplementary Figure 15.**
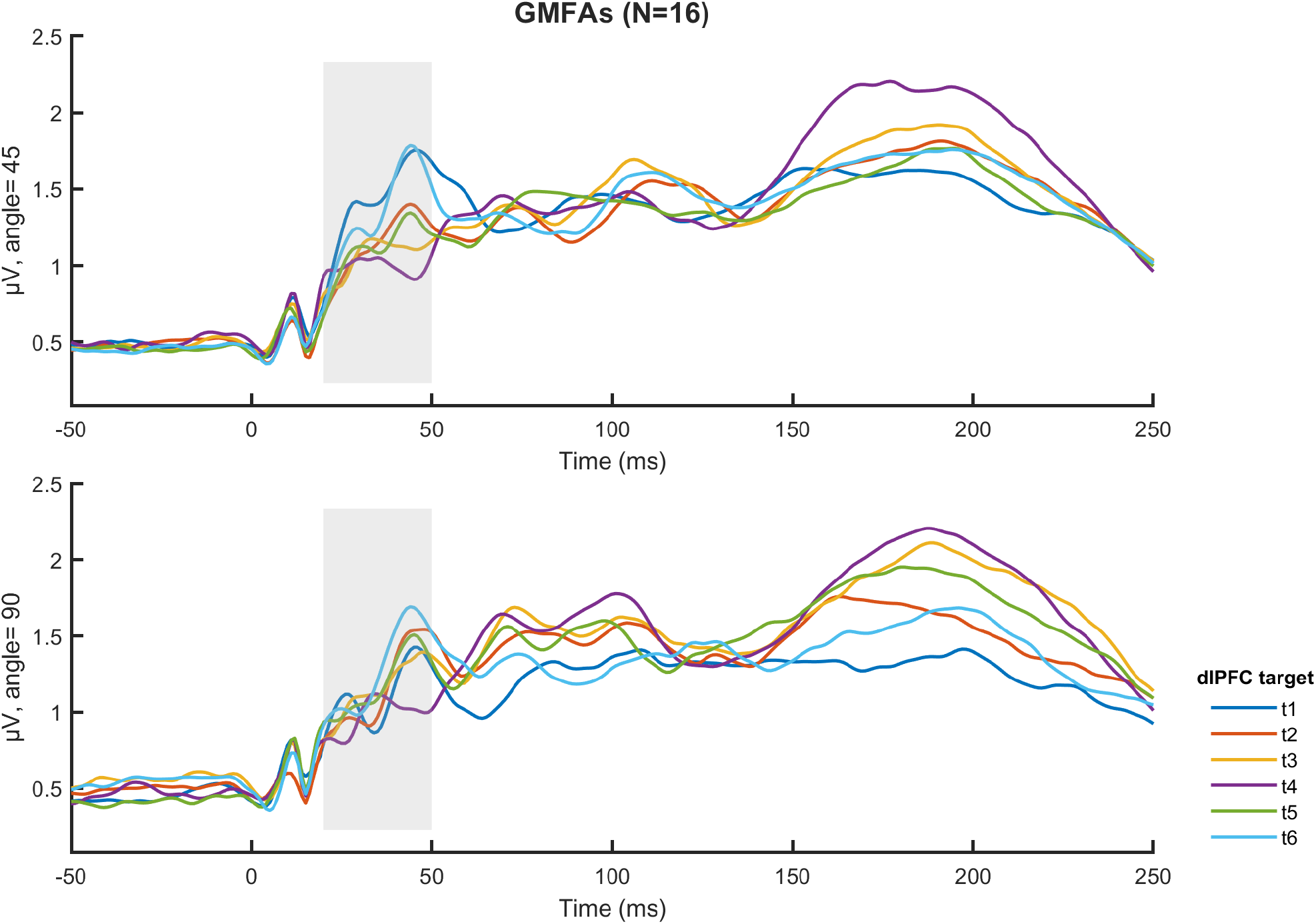
Mean GMFAs across targets (N=16). Upper panel represents 45 degrees coil angle and the lower panel represents 90 degrees coil angle

